# Glial cells promote infection by neurotropic Influenza A viruses *in vitro*

**DOI:** 10.1101/2025.06.20.660606

**Authors:** Nele Rieke, Maximilian Rohde, Melissa Käune, Lea Gabele, Mathias Müsken, Kristin Michaelsen-Preusse, Shirin Hosseini, Martin Korte, Christian Sieben

## Abstract

Although influenza A viruses (IAV) are notorious respiratory pathogens, some IAV strains can reach and replicate in the central nervous system (CNS) *in vivo*. In particular, highly pathogenic avian influenza viruses (HPAIV) pose a threat for future pandemics and are linked to greater neurotropic potential. Moreover, neurotropic IAV strains have shown a more significant impact on the onset and pathogenesis of neurodegenerative diseases (NDD). However, despite its clinical relevance, the dynamics and cellular tropism of IAV infection in the CNS are not well understood. In this study, we analyzed the replication of HPAIV H7N7 in vitro using a primary murine triple co-culture system comprising neurons, astrocytes, and microglia. We found that microglia become highly infected early on and induce a strong pro-inflammatory response before undergoing apoptosis. Using fluorescence microscopy with automated single-cell profiling, we found that, in contrast to non-neurotropic H3N2, the H7N7 nucleoprotein accumulated in neuronal somata throughout the infection without being transported into dendrites or axons. A combination of single-cell and co-culture replication assays led us to conclude that astrocytes are the primary virus producers of HPAIV H7N7 in the CNS. Our results illuminate the acute phase of neurotropic IAV infection, highlighting its implications for the association between IAV and the development of neurodegenerative diseases.

## Introduction

Influenza A viruses (IAVs) are a major human health threat that cause annual epidemics and sporadic pandemics. As one of the most devastating IAV pandemics, the “Spanish Flu” from 1918 has caused the death of approximately 50 million people, with one-third of the world’s population infected [1]. Interestingly, even though Influenza is a respiratory disease, the pandemic from 1918 was accompanied by a “mysterious sleeping disorder” named “Encephalitis lethargica” [2]. Decades after the Spanish flu, there was a rise in the incidence of Parkinson’s disease (PD), starting an ongoing debate about whether IAV infection could cause the onset and progression of neurodegenerative diseases [3, 4].

IAVs are enveloped negative-sense single-stranded RNA viruses carrying eight viral RNA segments. IAVs are transmitted between humans via aerosols and infect epithelial cells of the upper and lower respiratory tract. Although IAV infection typically leads to a respiratory disease, a wide range of neurological complications have been reported, including those affecting the central nervous system (CNS) [5]. In particular, highly pathogenic avian influenza viruses (HPAIV), including strains from the subtypes H5 and H7 can exert broad neurotropic potential in mammalian species, correlating with neurological disease severity [6, 7]. While the cellular tropism and details about the IAV infection mechanism have been established for cells derived from different tissues outside the CNS, little is known about IAV tropism and replication in neuronal cell types.

Neurotropic IAV strains such as HPAIV can reach the brain via different cranial nerve endings or hematogenous routes [8–10]. Following infection with the neurotropic IAV strain H7N7, a decrease in postsynaptic terminals was observed *in vivo* [11], and was later refined to have a sex-specific character *in vitro* [12]. Spine loss *in vivo* was further associated with microglia activation, cognitive impairments and an inflammatory transcriptome signature involving neuron- and neuroglia-specific genes [11]. Although accumulating evidence suggests that neuroinflammation is highly involved in the development of neurodegenerative diseases, including PD [3, 13], the mechanistic details about neurotropic IAV infection still remain elusive.

Here, we investigate the infection and replication kinetics of neurotropic IAV strains in murine primary neuronal cells, as well as the cellular reactions to infection, to better understand neurotropism and its role in the development of neurodegenerative diseases. We found that neurons and glial cells (microglia and astrocytes) become infected rapidly. While microglia undergo apoptosis during the first 24 h post-infection, neurons do not show signs of progressing infection, leaving astrocytes as a likely driver of neurotropic IAV infection *in vivo*. Our results shed light on the acute phase of neurotropic IAV infection with implications for the association of IAV and the development of neurodegenerative diseases.

## Methods

### Virus production in cell lines

IAV recombinant mouse-adapted A/Seal/Mass/1/80 (rSC35M; H7N7; kindly provided by Prof. Dr. Klaus Schughart, HZI; [14]), and rSC35M-NS1-GFP (kindly provided by Dr. Peter Reuther; Universitätsklinikum Freiburg; [15]) were propagated in Madin-Darby Canine Kidney (MDCK) cells. MDCK cells were cultured in DMEM (Life Technologies, 52100021) containing 10 % fetal bovine serum (Sigma-Aldrich, F7524). For propagation of the virus, MDCK cells were used at a confluency of around 90 %. The cells were washed three times with warm dPBS (Gibco, 10010-015) and the infection medium containing DMEM supplemented with 0.2 % bovine serum albumin (BSA) (Sigma-Aldrich, A7409), 1 % Penicillin/Streptomycin (Gibco, 15070-063) and 2.5 ng/ml N-Acetyl-Thiazolium-Bromid (NAT) (Sigma-Aldrich, T6763) was added together with an appropriate amount of virus. After 1 h at 37 °C, 5 % CO_2_ the infection medium was removed, the cells were washed once with warm dPBS supplemented with 0.2 % BSA and the medium was changed to fresh infection medium. The cells were incubated for around 40 h at 37 °C, 5 % CO_2_ until the observation of cytopathic effects. The supernatant was harvested and centrifuged for 10 minutes at 4 °C, 1500 rpm, transferred into a new falcon tube on ice, aliquoted into appropriate volumes and stored at - 80 °C until further usage.

### Virus production in embryonated chicken eggs

IAV mouse-adapted A/Hong-Kong/1/68 68 (maHK68; H3N2; kindly provided by Prof. Dr. Klaus Schughart, HZI; [16]) and H5N1 clade 2.3.4.4b (A/black-headed Gull/Ger-BW/AI01419/2023; kindly provided by Friedrich-Löffler Institute) were produced in embryonated chicken eggs. Specific pathogen-free chicken eggs were bred for 10 days at 37 °C, 50-70 % humidity while being rotated regularly. For the infection, the virus was diluted 1:1000 in PBS in 15 ml Falcon tubes on ice. Blunt ends of eggs were disinfected using iodine and a small hole was pierced into the eggs. With a syringe, 200 µl of the virus dilution was injected into the allantoic fluid. The injection side was sealed using glue and the infected eggs were incubated for 48 h at 37 °C, 50-70 % humidity. Afterwards, the eggs were incubated overnight at 4 °C. To harvest the virus, eggs were opened using a knife and outer membranes were removed. The allantoic fluid was extracted using a pipette and transferred into a Falcon tube and stored on ice. Aliquots were made and stored at -80 °C.

### Determination of the infectious virus titer

To determine the titer as FOCI-forming units (FFU), FOCI assays were performed. For this, 3*10^4^ MDCK cells/well were plated in a 96-well plate with flat bottom and incubated for 24 h at 37 °C, 5 % CO_2_. Next, virus dilutions were prepared in infection medium containing DMEM with 0.1 % BSA (Sigma-Aldrich, A7409) and 2.5 ng/ml NAT. The MDCK cells were washed with 100 µl infection medium before 50 µl of the virus dilution was added to each well. After 1 h at 37 °C, 5 % CO_2_, the infection medium was discarded and cellulose-DMEM was added containing 50 % 2xDMEM (2.68 % DMEM (Life Technologies, 52100021), 0.75 % NaHCO_3_ (Carl Roth, 6885.1), 2 % Glutamax (Life Technologies, 35050061), 2 % penicillin/streptomycin, adjusted with HCl (Carl Roth, 9277.1) to pH 7.2 and sterile-filtered using 0.2 μm unit (Sarstedt, 83.1826.001), 1 % carboxymethylcellulose sodium (Sigma-Aldrich, C9481), 0.1 % BSA and 2.5 ng/ml NAT in H_2_O. The cells were incubated for 24 h at 37 °C, 5 % CO_2_. After the overlay was tapped out carefully the cells were washed twice with 100 µl dPBS and fixated with cold 100 % ethanol (J.T. Baker, 8025) for 10 minutes. The experiment was performed at room temperature. Next, the cells got washed twice with dPBS and Quencher (1.5 % glycine (Serva, 23391.03) and 0.5 % Triton X100 (Sigma-Aldrich, T8787) in PBS) was added. After 10 minutes incubating the cells got washed once with washing buffer containing 0.5 % Tween 20 (Sigma-Aldrich, 8.22184) in PBS. The primary antibody (Virostat, 1301-1) was diluted 1:1000 in blocking buffer consisting of 0.5 % Tween 20 and 1 % BSA in PBS and added to the cells for 1 h. Afterwards, the cells were washed three times with washing buffer and the secondary antibody (KPL, 5220-0362) diluted 1:1000 in blocking buffer was added for 1 h. The cells were washed three times with washing buffer. Last, the peroxidase substrate (KPL, 5510-0030) was added for 15-30 minutes and the plate was rinsed with water. FOCI were counted and the titer was calculated as FFU per ml.

### Cell lines

MDCK and H4 cells were cultured in DMEM (Gibco, 6195-026) supplemented with 10% FCS (Capricorn Scientific, FCS-62A) in cell culture flasks (Sarstedt, 83.3911). Cells were detached using trypsin/EDTA solution (1X) (Sigma, T3924) and a portion of cells was stained with tryphan blue solution (Gibco™, 15250061) and counted using a Neubauer chamber (Brand™, 717805).

### Primary murine microglia monocultures

All protocols for providing cultures obtaining from C57BL6J and CX3CR1GFP+ mice in this project were reviewed and approved by the local committees at TU Braunschweig and the relevant authorities (LAVES, Oldenburg, Germany §4 (09/22) TSB TU BS), in accordance with Germany’s national Animal Welfare Act (‘Tierschutzgesetz’ in the version published on May 18, 2006 (BGBl. I S. 1206, 1313)).

Mouse C57BL/6 P3-5 pubs were decapitated and brains were transferred into cold 1xHBSS (Gibco, 14185-045). Hippocampi were extracted, transferred into 15 ml tubes and centrifugated for 15 minutes at 400g at 4 °C. Supernatant was removed and 5 ml cold HBSS was added. Tissue was resuspended by pipetting followed by filtering into a 50 ml tubes using 100 µm filters (Greiner, 542000) that were moistened with HBSS. Afterwards, the filter was rinsed with 5 mL HBSS and the suspension was centrifugated for 5 minutes at 400g at 4 °C. Supernatant was removed and pellet resuspended in 1 ml cell culture medium per brain using 10 ml pipettes. 1 ml cells were transferred to pre-coated 75 cm^2^ cell culture flasks. For coating, the flasks (Greiner, 658170) were covered during the procedure with 2 ml of undiluted Poly-D-lysine (Sigma, 6407) und incubated for 5-30 minutes on a shaker. Afterwards, the flasks were rinsed twice with PBS, filled with 10 ml DMEM (Capricorn, CP23-6230) +10 % FCS (Capricorn, 12B) + P/S (Gibco, 15070-063) and stored at 37°C, 10 % CO_2_ until usage. Cells were incubated for 17-21 days. At day 3 100 % of cell medium was exchanged. Afterwards, every 7 days 10 ml cell medium was exchanged. After reaching confluency, the cell culture flasks were shaken at 220 U/min at 37 °C for 2 h. Supernatants contain microglia, while astrocytes were still attached to the bottom. Supernatants were transferred into 50 ml falcons and centrifugated for 5 minutes at 400g at room temperature. Supernatants were removed and pellets resuspended in 1 ml DMEM with 10 % FCS. Cells were counted using a Neubauer chamber and plated on uncoated coverslips. They were incubated at 37°C 5 % CO_2_ for 24-48 hours and then used for experiments.

### Primary murine astrocyte monocultures

Primary murine astrocyte cultures were prepared as described by Lonnemann et al. [17]. Briefly, neonatal mouse brains (P3-4) were used. After removal of the brain, the hippocampus and meninges were carefully removed. The cortices were transferred on ice to fresh HBSS 1 X, and the tissue was homogenized using a 10 ml pipette. The HBSS was washed off and the brains were transferred into a dissociation solution containing DNAse for 30 minutes at 37 °C, and the remaining tissue pieces were further dissociated by pipetting. After centrifugation at 800 rpm for 7 minutes, the supernatant was removed and the cells were resuspended in culture medium (DMEM supplemented with 10% FCS, 1% penicillin/streptomycin) and placed on a 40 µm cell strainer and finally transferred to a 75 cm^2^ cell culture flask pre-coated with poly-D-lysine. After reaching confluency, the cells were shaken overnight at 220 rpm to remove other glial cells, and then astrocytes were passaged. The cells were passaged three times until the experiments were performed.

### Poly-L-Lysine Coating of Coverslips

Glass coverslips with a diameter of 13 mm and a thickness of 1 mm (VRN, 631-1578) were incubated in 10 M NaOH for 3–5 h at 100 °C. The coverslips were washed five times with distilled water for 20 min and subsequently sterilized at 225 °C for 6 h. After cooling, the coverslips were coated with 0.5 mg/ml poly-L-lysine (Sigma-Aldrich, P2636) dissolved in boric acid buffer at 37°C for 2-3 h and washed five times with distilled water. After drying, they were stored at 4°C until further use.

### Triple co-culture and co-culture preparation containing neurons, astrocytes and microglia

Primary embryonic hippocampal cultures were prepared as previously [12, 18] described. In brief, C57BL/6J mice were decapitated at embryonic day 17.5 (E17.5) and embryos were divided by sex [19]. Both hippocampi were isolated and enzymatically dissociated in 1X trypsin/EDTA solution (Sigma-Aldrich, T3924) for 25 minutes at 37 °C. The tissue was then mechanically homogenized using a narrowed Pasteur pipette. Homogenized hippocampi were centrifuged for 5 minutes at 1500 rpm. The cell pellet was resuspended in DMEM medium (Gibco, 6195-026) supplemented with 10 % FCS (Capricorn-scientific, FCS-62A). This was followed by three washes in cell culture medium. As cell culture medium neurobasal medium (Gibco, 21103-049) supplemented with N2 (autoclaved), B27 (Invitrogen, 17504-001) and 0.5 mM L-glutamine (Invitrogen, 25030-024) was used to obtain co-cultures of primary neurons and astrocytes. 7*10^4^ cells per well were plated onto poly-L-lysine-coated 13 mm diameter coverslips (described above) in a 24-well plate (Sarstedt, 83.3922) and cultured for 21 days in cell culture medium. Co-cultures were maintained in an incubator at 37 °C, 5 % CO_2_ and 99 % humidity, with 20 % of the cell medium changed once a week.

To obtain the triple co-culture consisting out of neurons, astrocytes and microglia, 3* 10^4^ microglia, prepared as described above, per well were plated on the embryonic hippocampal culture. Triple co-cultures were stored in an incubator at 37 °C, 5 % CO_2_ and 99 % humidity and treated 72 h after microglia plating.

### Infection of primary cell culture

The cell culture medium was collected. Next, 200 µl infection mix consisting of the appropriate amount of virus or PBS diluted in fresh NB^+^ medium was added to the neuronal cells. After 1 h at 37 °C, 5 % CO_2_ the infection mix was removed and cells were washed three times with 200 µl pre-warmed fresh NB^+^ medium to get rid of unbound virus particles in the medium. The post-infection medium contained 80 % of the collected cell culture medium and 20 % fresh NB^+^ medium. Per well, 500 µl post-infection medium was added and cells were incubated at 37 °C, 5 % CO_2_. Cells were fixed with 4 % PFA in PBS.

### Infection of cell lines

The cell culture medium was collected and cells washed once with infection medium, being DMEM with 0,1 % BSA and 2.5 ng/ml NAT. Next, 200 µl infection mix consisting of the appropriate amount of virus or PBS diluted in infection medium was added to the neuronal cells. After 1 h at 37 °C, 5 % CO_2_ the infection mix was removed and cells were washed three times with 200 µl infection medium to get rid of unbound virus particles in the medium. Per well, 500 µl infection medium was added and cells were incubated at 37 °C, 5 % CO_2_. Cells were fixed with 4 % PFA in PBS.

### Immunocytochemistry

Fixed cells were incubated for 1h at RT with blocking buffer consisting out of PBS with Triton X100 and 0.1 % BSA. Primary antibodies were diluted in blocking buffer and added to the cells for either 1 h at RT or over night on a shaker at 4°C. Afterwards the cells were washed three times with PBS. Secondary antibodies were diluted in blocking buffer and added to the cells for 1 h at RT. A list of antibodies used in this study is shown in **Supplementary Table 1**. The cells were washed three times with PBS and Hoechst staining (Thermo Scientific, 33342) was added 1:10.000 for 5 minutes. Cells were washed once with PBS for 5 minutes. For the lectin stainings, the respective lectins were diluted in blocking buffer and added for 5 minutes. The cells were washed once with PBS afterwards. Before being mounted the cells were washed with MilliQ water.

### Scanning electron microscopy

Scanning electron microscopy was performed as described previously [20]. In brief, triple co-cultures on cover slips were fixed with 5% formaldehyde and 2% glutaraldehyde (final concentration), washed twice in TE buffer (20 mM TRIS and 1 mM EDTA, at pH 6.9), and dehydrated on ice in a graded series of acetone (10%, 30%, 50%, 70% and 90%) for 10 min each step, followed by two steps in 100% acetone at room temperature. Critical point drying with liquid CO2 was performed with the CPD300 (Leica Microsystems) and sputter coating (55 s at 45 mA) with the SCD500 (Bal-Tec, Liechtenstein) to coat the samples with a thin gold–palladium film. Image acquisition was performed with a field emission scanning electron microscope Zeiss Merlin (Zeiss, Oberkochen) using both, the Everhart Thornley HESE2 detector and the in-lens SE detector and with an acceleration voltage of 5 kV and operated with the SmartSEM (version 6.08, Carl Zeiss Microscopy Ltd) software.

### Fluorescence microscopy

Microscopy was performed using an EVOS™ M500™ (Invitrogen™). We used the same imaging properties within one dataset for quantification experiments. The images were acquired randomly and each channel of an image was saved as a separate TIF-file.

### Single-cell infection- and cell-type quantification in Python

Single-cell analysis of infected cell was performed in python (version 3.9). Each channel of an image was saved as a TIF-file. The channel containing nuclei was segmented using the tool StarDist [21] (StarDist version 0.9.1; CSBDeep version 0.8.0). For the other channels an individual threshold was determined. For this, the most frequent value (mode) of a channel was identified and multiplied with a specific value. This multiplier could be adapted if the result of the segmentation was not sufficient. Gaussian filter was applied (Scipy version 1.9.1) followed by thresholding and generation of a label image using scikit-image (version 0.19.2). After binarization each area in each channel was validated to ensure that cells that localize close to each other were detected as separate. For this, all nuclei that lie within the coordinates of a particular area were counted. If the area did not co-localize with a nucleus it was deleted. In contrast, if the area had more than exactly one nucleus, it became adjusted. For adjustment, a rectangle was set up around each area. Within this rectangle the mode was used as a threshold and the multiplier is automatically adjusted in a stepwise manner until each area contained exactly 1 nucleus. After segmentation the cells were classified. For classification, the code iterated through all nuclei of an image and checked for signal in the other channels that colocalized to a certain extend (>75% overlap). The algorithm initially tested if a nucleus overlapped with an area in the microglia channel (marker: Iba1). If there was colocalization, the nucleus was classified as a microglia. If there was no overlap the algorithm tested for colocalization between the same nucleus and an area in the neuronal channel (marker: NeuN). Again, overlap with a channel classified this nucleus as the particular cell type. If there was no overlap with microglia and neurons, a nucleus was classified as an astrocyte.

For detection of infected and non-infected cells, the nuclei were tested for colocalization with an area in the IAV channel (marker: NP). The classification results were combined into pandas DataFrames and exported CSV-files. Pie plots were generated using matplotlib (version 3.5.2). The segmentation result was saved for each channel individually in a separate folder as JPEG using matplotlib. Data handling was performed using Pandas (version 1.4.4) and numpy (version 1.21.5) library. The code is available at https://github.com/christian-7/Rieke_et_al.

### Quantification of virus export in primary cell culture

Based on the quantification data obtained by the script described above we compared the area sizes within the NP channel with the area sizes of the corresponding nuclei. For this we developed a script. When applying a threshold of more than 1,5×10^3^ pixel in the NP channel and less than 0.5×10^3^ pixel in the DAPI channel at the same time a cell was assumed to show NP export. The code is available upon request.

### Apoptosis assay of primary cell culture

The cells were infected as described before. For live cell analysis either a Caspase-3/7 dye (Sartorius, 4704) or a cell death marker (Sartorius, 4632) were used. After the cells were washed three times at 1 hour post infection (hpi) the live imaging dye was added at a 1:10^3^ dilution to the medium. The cells were placed in in an Incucyte® SX5 (Sartorius) at 37°C, 7.5% CO_2_ and measured at the setting described in supplementary table 2.

### Cell motility analysis

A Python script (version 3.10) was developed to analyze microglia movement based on image data. Image processing and cell tracking were performed using several functions. First, skimage (version 0.6.4) was used to identify related objects in the individual images of a GIF. The optimized_mg_detection function converted each frame into a binary image and filtered objects based on their area (250-5000 pixels). The cells were then labeled with measure.label and their properties extracted. Trackpy (version 0.6.4) was used to track the microglial cells across multiple frames. Cells with similar positions in consecutive frames were linked using a search radius of 35 pixels and a memory of 5 frames. Each cell was assigned a unique ID. The cell movements were visualized using Matplotlib (version 3.9.2). The trajectories and bounding boxes of the cells were displayed graphically. Finally, the movement data and cell areas were saved in a table. This contains information on distances, cell areas and average values per individual image. The code is available upon request.

### Live-cell microscopy analysis

A Python script (version 3.10) was developed for the quantitative analysis of cell populations. Image processing included conversion to grayscale and segmentation by dynamic thresholding based on the pixel intensity mode, i.e. the most frequent pixel value in the image. Numpy (version 1.26.4) was used to calculate the intensity mode and determine the threshold values. Segmentation was performed using Scikit-Image (version 0.24.0) and OpenCV (version 4.10.0), with OpenCV responsible for threshold application and image processing. Objects below a minimum area of 250 pixel² (CX3CR1-labeled microglia) or 50 pixel² (caspase-3/7-positive cells) were excluded. Thresholds were calculated individually for each image acquisition based on the pixel intensity mode: 2.5-fold mode for CX3CR1-labeled microglia and 4-fold mode for caspase-3/7-positive cells. Data processing and calculations were carried out with Pandas (version 2.2.3), while Matplotlib (version 3.9.2) was used for graphical representation. Image processing and formatting of the output images was performed with Pillow (version 10.4.0). The algorithm classified cells into three categories: CX3CR1^+^caspase-3/7^-^ cells (green), CX3CR1^+^caspase-3/7^+^ cells (green and red), and CX3CR1^-^caspase-3/7^+^ cells (initial green, later red). In the cytotox assay, classification into cytotox-positive and cytotox-negative cells was performed. The code is available upon request.

### Multiplex cytokine assay

Supernatants from infected neuronal cells were collected and stored at -80 °C as described above. Cytokines were measured by LEGENDplex™ Mouse Inflammation Panel (13-plex) with Filter Plate (Biolegend, San Diego, CA, USA, #740150) according to the manufacturer’s protocol using BD™ LSR-II SORP.

### Statistical analysis

Statistical analysis was performed in Graphpad/Prism (version 10.1.1.). Normally distributed data was analyzed by one- or two-way ANOVA. For multiple comparison analysis, ANOVA was followed by Tukey’s multiple comparison test, otherwise an uncorrected Fisher’s LSD test was performed. The data was tested for normal distribution by D’Agostino-Pearson omnibus, Anderson-Darling, Shapiro-Wilk and Kolmogorov-Smirnov tests. If data did not pass normality testing, a Kruskal-Wallis test with Dunn’s multiple comparison test was performed. For statistical analysis, data from male and female-derived animals as well as from mixed cell cultures was pooled. All experiments were evaluated in a strictly blind fashion.

## Results

### Infection with neurotropic H7N7 shows faster progression than non-neurotropic H3N2 in neuronal cells

To investigate the cellular tropism and infection progression of neurotropic H7N7, we infected a murine triple co-culture with 8×10^5^ FFU corresponding to a multiplicity of infection (MOI) of 1 FFU/cell for 6 and 24 h. In addition to the neurotropic IAV strain (rSC35M, mouse-adapted A/Seal/Mass/1/80 (H7N7)), we included a non-neurotropic mouse-adapted A/Hong-Kong/1/68 human IAV strain (maHK68 (H3N2)), for comparison in our studies. To obtain a structural overview of our triple co-culture system, we performed scanning electron microscopy, which highlighted the extensive network between the cells in our cell culture system (**Supplementary Figure 1**). To leave the cellular network intact during our analysis, we thus decided to develop a microscopy-based readout for determining cell type and infection status. Infected cultures were fixed at different time points and stained using antibodies directed against cellular and viral markers as described in the methods section. Multi-color fluorescence microscopy images were taken using low-magnification (10x) wide-field microscopy and the images were analyzed using a custom Python script. The general workflow is depicted in **Figure 1**.

**Figure 1:**
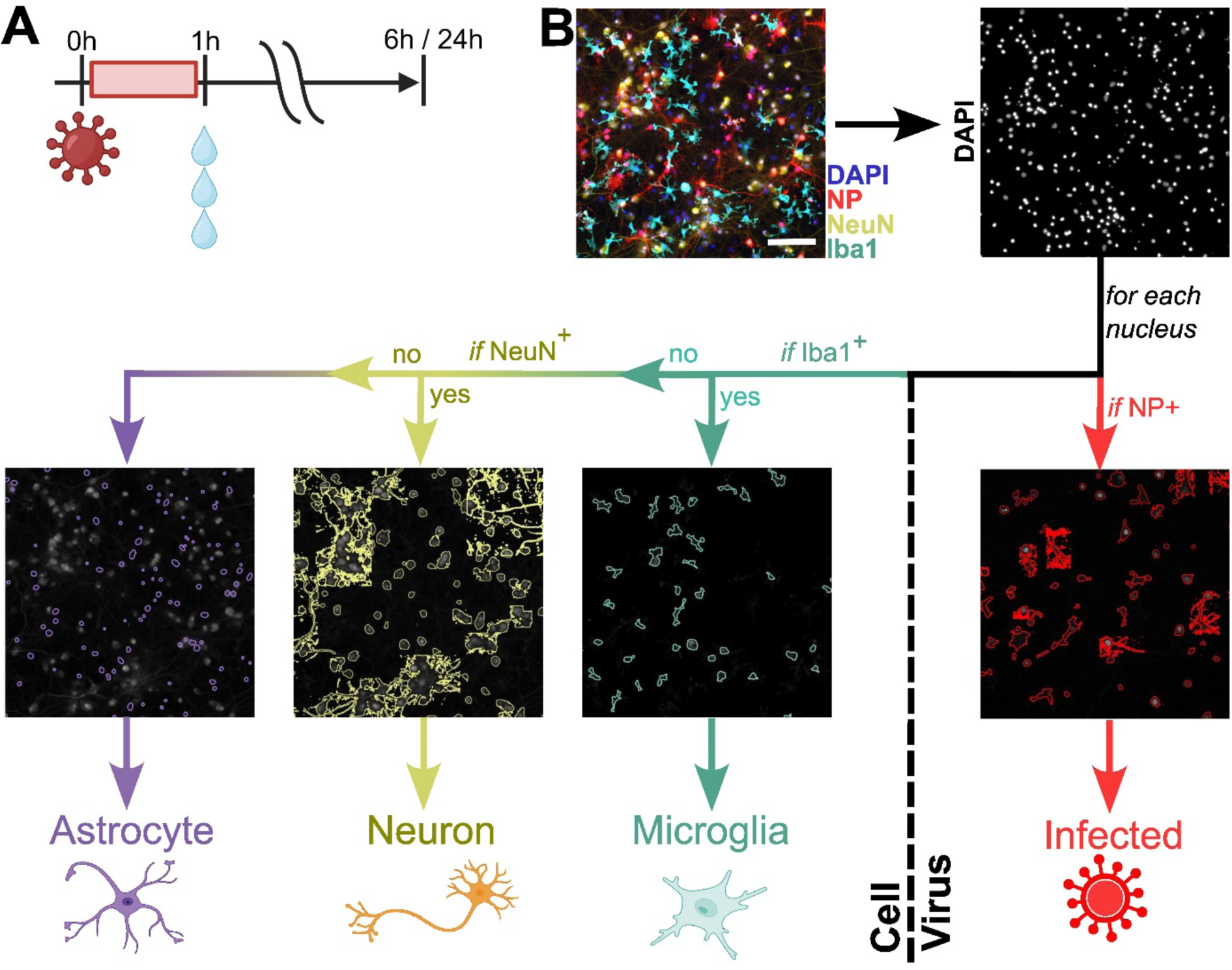
Single-cell profiling pipeline for the classification of cell type and infection status. **A**) Cells were infected with IAV and incubated for 1h. Afterwards, cells were washed three times to remove unbound viruses. Fixation and extraction of supernatants was performed at 6 and 24hpi. **B**) We developed a computational pipeline to automatically analyze all images within a specified folder. Input are the individual channels of an image in TIF-format. First, the channel containing stained nuclei was opened und nuclei were segmented using StarDist [21]. Next, all other channels were segmented using the most frequent value in that image multiplied by an adaptable value as threshold. Cell type classification was performed as followed: For each nucleus, the algorithm initially checked for colocalization (>75% overlap) with any area in the Iba1 channel. If there were overlapping areas, this nucleus was classified as microglia, before colocalization with the NeuN channel was checked, if the nucleus was not a microglia. If the nucleus was Iba1- and NeuN-negative, the cell was classified as astrocyte. After cell type classification, the same nuclei were analyzed for colocalization with the IAV NP channel. Overlap with an area in the NP channel classified a nucleus as infected. The decision tree and respective classification results are shown. In the following, we use a color code with cyan for microglia, yellow for neurons and purple for astrocytes.

First, we compared the infection rates of the neurotropic H7N7 with the non-neurotropic H3N2 strain at 6 and 24 hpi (**Figure 2A**). The infection rate corresponds to the percentage of infected cells within the total cell population. Following H7N7 infection, we detected 22.3 % infected cells after 6 h compared to H3N2 with 7,9 % infected cells (P_6h_H7N7-H3N2_<0.0001). At 24 hpi there was no statistically significant difference between both strains with 17.1 % infected cells after H7N7 infection compared to 11.9 % for H3N2 (P_24h_H7N7-H3N2_=0.144). Neither strain showed a statistically significant difference in the amount of infected cells between 6 and 24 hpi (P_H7N7_6h-24h_= 0,0757; P_H3N2_6h-24h_= 0,1486). To then check if H7N7 and H3N2 were able to replicate, we next analyzed the virus titer in the supernatant of infected cells via FOCI assay (**Figure 2B**). The virus titer following H7N7 infection was higher at 6 hpi with 2.21*10^3^ FFU/mL compared to 7.06×10^2^ FFU/mL at 24 hpi (P_H7N7_6h-24h_=0.0276). H3N2 showed lower viral titer at 6 hpi with 4.33×10^2^ FFU/mL (P_6h_H7N7-H3N2_=0.0001), which did not increase over time (P_H3N2_6h-24h_>0.9999). Together, these findings indicate that the neurotropic H7N7 can infect and replicate successfully in our triple co-culture system while the H3N2 infection might be effectively controlled.

**Figure 2:**
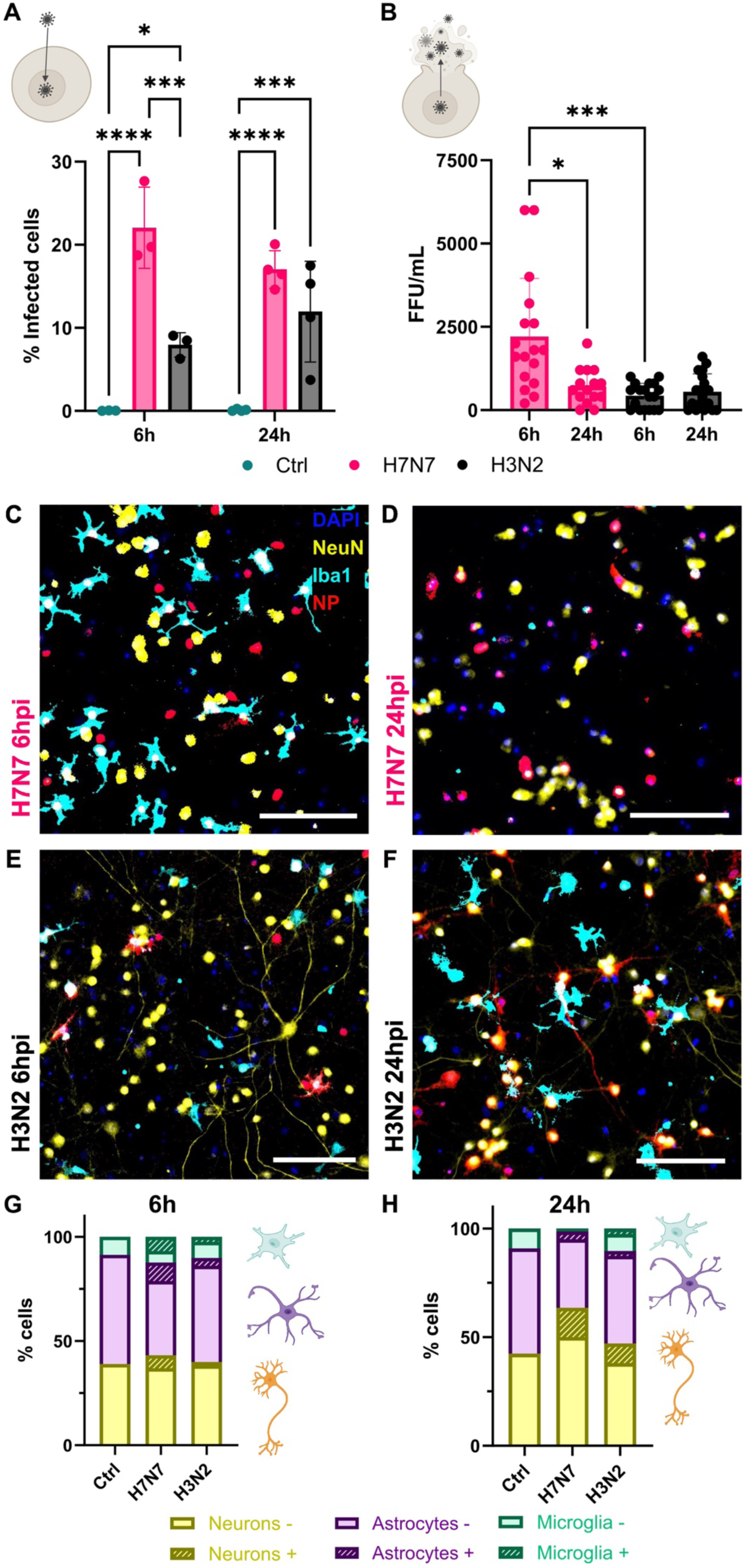
Infection with neurotropic H7N7 shows faster progression than non-neurotropic H3N2 in neuronal cells. **A**) Infection rate (in % NP^+^ cells) of both strains and control (mock infected) shows H7N7 exerts a faster infection progression. Significance calculated using two-way ANOVA followed by Tuke’s post hoc test. 6 hpi: N=3; 24 hpi: N=4 **B**) Replication rate (in FFU/ml) of both strains. Supernatants of infected cells were analyzed via foci assay and indicate H7N7 replication peaks at 6 hpi. Significance calculated using Kruskal-Wallis test with Dunn’s multiple comparisons test. N=4 **C-F**) Representative images of infected cells at 6 and 24 hpi. Blue: DAPI, yellow: NeuN, cyan: Iba1, red: NP. Scale bar: 200 µm. **G-H**) Cell composition in controls, H7N7 or H3N2 infected cells at 6 and 24 hpi. Dashed areas indicate infected cells. 6 hpi: N=3; 24 hpi: N=4. Statistical significances indicated as *p<0.05, **p<0.01, ***p<0.001, ****p<0.0001.

### H7N7 infection drastically changes the cell composition in the triple co-culture

We next investigated the contribution of each cell type to the progression of virus infection. Our custom analysis script performs automated cell segmentation and classification based on the signals present in the different fluorescence channels. A low fraction of oligodendrocytes is assumed to co-exist in our primary culture system but was not further considered here since to our knowledge, there is no report of infected oligodendrocytes *in vivo*. We highlight that the cell classification was based on the presence of neuronal or microglial markers solely and that all cells that were absent of either of them were classified as astrocytes which represent the vast majority in this population. With this analysis tool, the triple co-culture was calculated to be composed of 10.1 % microglia, 44.2 % astrocytes and 45.7 % neurons in control samples at 24 hpi. These proportions are in agreement with previous estimates that mimic the physiological state of the brain. An analysis of several *in vivo* studies revealed a cell count of 57.3 % neurons together with interneurons, 34.8 % astrocytes and 7.9 % microglia in the mouse hippocampus with a high variation in between different studies [22]. Of note, lectin staining revealed that neurons and microglia display sialylated glycans on their surface, while we could not detect any lectin signals on astrocytes (**Supplementary Figure 2**).

Representative images of H7N7- and H3N2-infected cells at 6 and 24 hpi are shown in **Figure 2C-F**. The corresponding cell composition is depicted below in **Figure 2G-H** and statistical analysis is shown in **Supplementary Figure 3**. Interestingly, we found that infection with H7N7 induced a strong decrease of the total microglia proportion from 12.4 % at 6 hpi to 1.4 % at 24 hpi, corresponding to a reduction by 88.6 % cells (P_H7N7_6h-24h_=0.0011). In contrast, H3N2 induced an increase in microglia abundance by 29.2% that was not statistically significant (P_H3N2_6h-24h_=0.12), similar to control samples treated with PBS that induced a 34.1% microglia increase (P_Ctrl_6h-24h_=0.5041). Due to the almost complete loss of microglia after H7N7 infection, the relative amount of neurons increased from 43.1 % at 6 hpi to 63.5 % at 24 hpi, corresponding to an increase by 47.4 % cells. Astrocytes decreased upon H7N7 infection by 21.1%, but the differences were not statistically significant (P_H7N7_6h-24h_=0.2726). Neither H3N2 nor PBS treatment induced any statistically significant cell number changes of neurons, astrocytes or microglia.

### Glia-to-neuron switch in the IAV cell tropism during infection progression

To further investigate the drastic changes in cellular composition upon IAV infection, we investigated the cell tropism of both IAV subtypes (**Figure 2G-H**). Concerning neurons, we observed IAV strain- and time-dependent effects on the infection status. For H7N7 infection, there was an increase from 14.0 % of neurons being infected at 6 hpi compared to 21.9 % at 24 hpi (p_H7N7_6h-24h_=0.0591). This increase was more pronounced after H3N2 infection from 4.2 % at 6 hpi compared to 19.1 % at 24 hpi (P_H3N2_6h-24h_ =0.0013). We did not find significant differences between H7N7 and H3N2 infection at 6 and 24 hpi (P_6hpi_H7N7-H3N2_=0.0743; P_24hpi_H7N7-H3N2_ = 0.7715) (**Supplementary Figure 3B**).

Next, we analyzed the amount of infected astrocytes and found strain- and time-dependent variations. At 6 hpi, more astrocytes were infected with H7N7 (20.0 %) compared to 7.4 % after H3N2 infection (P_6hpi_H7N7-H3N2_=0.0003). After 24 h, there was no significant difference with 10.5 % of astrocytes being infected with H7N7 and 6.6 % with H3N2 (P_24hpi_H7N7-H3N2_=0.1219). For H7N7, but not for H3N2, the amount of infected astrocytes decreased from 6 to 24 hpi (P_H7N7_6h-24h_=0.0003; P_H3N2_6h-24h_=0.3095) (**Supplementary Figure 3E**).

We observed significant differences in the amount of infected microglia between H7N7 and H3N2 as well as in the kinetics of infection. At 6 hpi, the infection was more pronounced after H7N7 infection with 56.9 % of microglia being infected compared to 25.6 % after H3N2 infection (P_6h_H7N7-H3N2_=0.002). Afterwards, the H7N7-infected microglia population strongly decreased as described above. At 24 hpi there were no statistical differences in the amount of microglia that were infected between both strains (Mean_H7N7_24hpi_ = 19.3 %; Mean_H3N2_24hpi_ = 12.2 %; P_24hpi_H7N7-H3N2_ = 0.5251; P_H7N7_6-24hpi_ <0.0001; P_H3N2_6-24hpi_ =0.072) (**Supplementary Figure 3H).**

Since H7N7 induced an overall higher infection rate compared to H3N2 (**Figure 2A**) and because the cell culture composition changed during the course of infection, we further calculated the composition of the infected population (**Figure 2G-H**, **Supplementary Figure 3C, F, I**). The H7N7-infected population at 6hpi was composed of 27.0 % neurons, 42.7 % astrocytes and 30.4 % microglia. At 24 hpi, it was composed of 76.9 % neurons, 20.1 % astrocytes and 0.4 % microglia. For H3N2, the infected population at 6 hpi was composed of 20.8 % neurons, 46.3 % astrocytes and 33.0 % microglia and changed to 70.4 % neurons, 16.1 % astrocytes and 5.1 % microglia at 24 hpi.

Taken together, at 6hpi, astrocytes showed the number of infected cells while microglia exhibited the highest infected fraction per cell type, regardless of the IAV subtype. At 24hpi, the glial cells showed decreased numbers among infected cells as well as among infected cells per cell type, whereas neurons displayed an increase in both ratios. This indicates that microglia contributed more significantly to early infection for both strains, while neurons exhibited higher infection rates in later stages. Although astrocytes showed a stronger contribution early on, they did not experience the same level of cellular disappearance as was observed for microglia.

### Neurotropic potential of HPAIV H5N1 clade 2.3.4.4b in vitro

Due to the observed strong differences between H7N7 and H3N2, we wanted to test the cellular tropism of HPAIV H5N1 in our murine triple co-culture. For this, we used a reduced virus concentration to investigate the replication efficacy in CNS-derived cells between the two HPAIV strains and the seasonal H3N2 IAV. To this end, triple co-cultures were infected with 2×10^4^ FFU/ml of HPAIV H5N1 clade 2.3.4.4b, H7N7 and H3N2 corresponding to MOI = 0.05, assuming a total of 4×10^5^ cells/ml. Despite not being mouse-adapted, avian H5N1 was able to infect all cell types in the murine triple co-culture system as shown in **Figure 3A**, but we observed differences in the infection rates between the tested IAV strains as shown in **Figure 3B**. H7N7 resulted in the highest percentage of infected cells, with 10.7% at 6 hpi, compared to H5N1 at 1.8% and H3N2 at 0.3%. This trend was also detected at 24 hpi, where H7N7 showed the highest fraction of infected cells with 19.6 % compared to H5N1 with 10.4 % and H3N2 with 2.9 % (P_H7N7-H3N2_ = 0.0018; P_H7N7-H5N1_ = 0.124). When comparing the amount of infected cells after infection with MOI = 0.05 and MOI = 1, we found that, at 6 hpi, the differences in the infection rates were statistically significant for both, H7N7 and H3N2 (P_H7N7_ < 0.0001, P_H3N2_ =0,0027). In contrast, at 24 hpi, the amount of infected cells was not statistically significant between both virus concentrations for H7N7 any more (P_H7N7_ =0.5057, P_H3N2_ =0,0269, not shown). We therefore conclude that the H7N7 might have reached the maximum infection capacity at MOI = 1 already at 6hpi and at MOI = 0,05 at around 24hpi.

**Figure 3:**
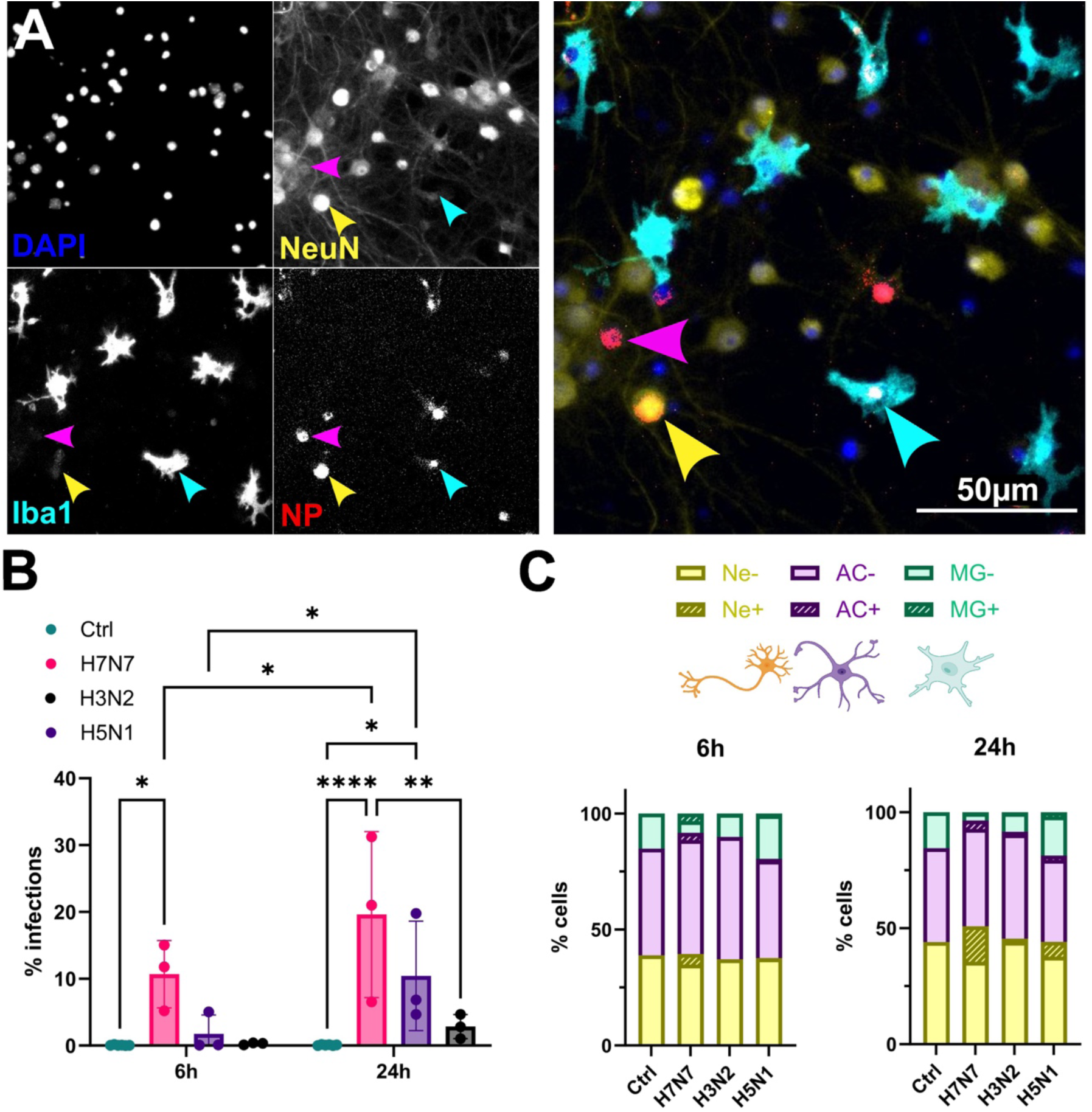
Neurotropic potential of HPAIV H5N1 clade 2.3.4.4b *in vitro*. **A**) Representative images of H5N1 infected cells at 24 hpi. Blue: DAPI, yellow: NeuN, cyan: Iba1, red: NP. Colored arrowheads show infected cells of each particular cell type. **B**) The infection rate (in % NP+ cells) shows H7N7 exerts a faster and stronger infection compared to avian H5N1 and mouse-adapted H3N2. Significance calculated using two-way ANOVA followed by Tuke’s post hoc test. **C**) Cell composition in H7N7, H5N1 or H3N2 infected cells at 6 or 24 hpi. Dashed areas indicate infected cells. Statistical significances indicated as *p<0.05, **p<0.01, ***p<0.001, ****p<0.0001. N=3.

The cell culture composition did not show statistically significant differences after infection with either virus at MOI = 0.05 (**Supplementary Figure 4 A, D, G**). However, there was a trend of microglia reduction after infection with H7N7 (**Supplementary Figure 4 G**).

We next investigated the cellular tropism of the different IAV strains and observed strain-dependent infection rates in all three cell types (**Figure 3C, Supplementary Figure 4**). At6 hpi, H7N7 showed the highest number of infected cells among neurons with 11.9 % in contrast to H5N1 with 0.3 % and H3N2 with 0.1 % (**Supplementary Figure 4B**). These differences were not statistically significant (P_H7N7-H5N1_=0.3248 P_H7N7-H3N2_=0.3084; P_H5N1-H3N2_>0.9999). After 24 hpi, the fraction of H7N7-infected neurons increased to 34.5 % (P_H7N7_6h-24h_ = 0.0026). The differences to H5N1 with 13.6 % and H3N2 with 3.4 % were also statistically significant (P_H3N2-H7N7_ =0.0006; P_H5N1-H7N7_ = 0.0224 ; P_H5N1-H3N2_ = 0.434) (**Supplementary Figure 4B**). We could also observe strain-dependent infection rates of astrocytes. At 6 hpi, H7N7 showed with 6.0 % the highest percentage of infected astrocytes compared to H3N2 with 0.1% (P=0.0248) and H5N1 with 1.7 % (P=0.1334) (**Supplementary Figure 4E**). At 24 hpi, there were no statistically significant differences between H7N7 with 7.3 % and H5N1 with 5.2 % or H3N2 with 2.2 % (p_H7N7-H5N1_=0.6795; p_H7N7-H3N2_ = 0.0579; p_H5N1-H3N2_ = 0.4083) (**Supplementary Figure 4E**). Concerning microglia, we found 40.0 % infected with H7N7 after 6 h compared to 5.1 % for H5N1 (P_H7N7-H5N1_<0.0001) and 2.5 % for H3N2 (P_H7N7-H3N2_<0.0001). As mentioned above, the fraction of H7N7-infected microglia went down to 9,6 % after 24h with no statistically significant differences between the three strains.

We then quantified the composition of the H7N7- and H5N1-infected cell population (**Supplementary Figure 4 C, F, I**). At 6 hpi, and especially for H7N7, there was a more even distribution of the different cell types compared to 24 hpi. For H5N1, the composition switched from 19.1 % neurons, 29.5 % astrocytes and 51.4 % microglia at 6 hpi to 60.3 % neurons, 19.8 % astrocytes and 19.9 % microglia at 24 hpi. The change of the infected cell composition was more severe after H7N7 infection with 39.7 % neurons, 31.9 % astrocytes and 28.4 % microglia at 6 hpi and 81.0 % neurons, 17.1 % astrocytes and 1.9 % microglia at 24 hpi. Taken together, at MOI = 0.05, astrocytes and especially microglia had a high contribution to the infection at early time points, while there was a shift towards neuronal infections at later stages of infection as we have described above for MOI = 1. Comparing the amount of infected cells after infection with MOI = 0.05 and MOI = 1 indicates HPAIV H7N7 rapidly reached its maximum infection capacity at both initial viral concentrations.

### Glia cells show signs of productive H7N7 and H3N2 replication

Following the characterization of the infection rates for each CNS cell type, we aimed to understand which cell type could act as virus producer. To this end, we infected cells with H7N7 or H3N2 with MOI = 1 and performed an image-based single-cell analysis of the intracellular localization of the viral NP throughout the first 24 h of infection. The IAV NP initially accumulates in the nucleus to form progeny vRNPs, the infection stage we used before to quantify the infection state of the cells. Later during viral replication, NP is exported to the cytoplasm and eventually reaches the plasma membrane to assemble into new virions. We thus visualized the viral replication cycle of both strains by staining the viral NP in each cell type. In particular, we then assessed NP export from the nucleus as a sign for ongoing viral replication (**Figure 4A**). vRNP nuclear export is a critical step and depends on the ras-related nuclear protein and its depletion results in the nuclear accumulation of NP and the inhibition of the IAV replication [23].

**Figure 4:**
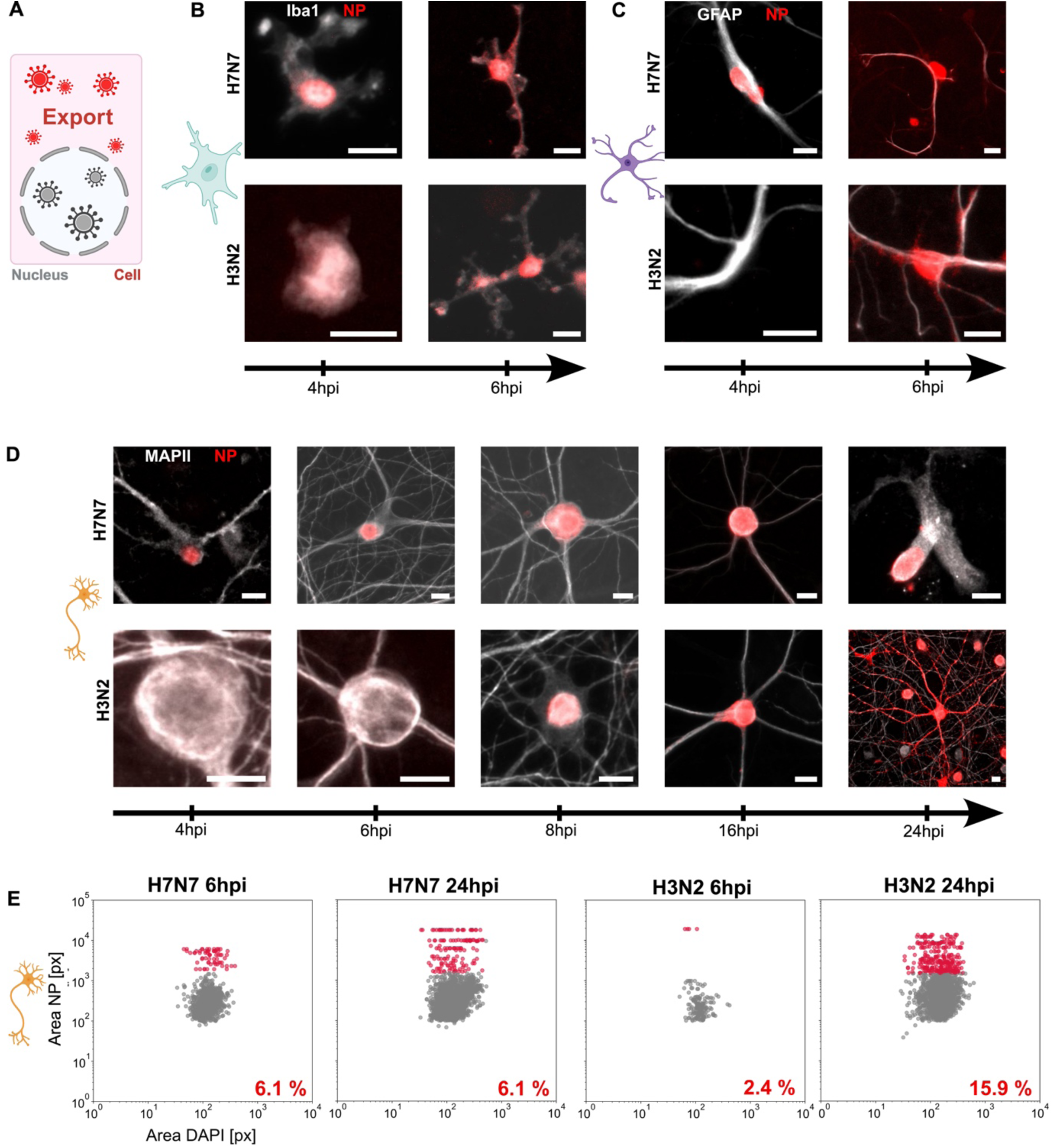
Single-cell analysis of the infection progression in neurons, astrocytes and microglia. **A**) Schematic representation of the expected export of viral NP from the nucleus into the cytoplasm of infected cells. **B**) Infection in microglia. H7N7 (upper panel) NP (red) localizes to the nuclei of H7N7-infected cells around 4 hpi and is exported at 6 hpi. For H3N2 (lower panel) the nuclear accumulation starts between 4 and 6 hpi and export is visible at 6 hpi. Scale bar 10 µm. **C**) Infection in astrocytes. NP localizes in the nuclei of infected cells around 4 hpi and is exported at 6 hpi. Scale bar 10 µm. **D**) Infection in neurons. H7N7 NP is found in the nuclei of infected cells as early as 4 hpi NP does not localize to any distal parts of neurons within 24 hpi. H3N2 NP becomes visible around 8 hpi and localizes to neurites starting around 16 hpi. Scale bar 10µm. **E**) Plots of NP^+^ cell area compared to the DAPI^+^ nuclear area at 6 hpi and 24 hpi after H7N7 or H3N2 infection. At 6 hpi two populations can be distinguished with a threshold of 1.2×10^3^ pixel per NP^+^ area. Assuming cells above the threshold show NP export, we detected 11.9% of infected neurons export NP at 6 hpi whereas 6.5 % of infected neurons export NP at 24 hpi. For H3N2 we detected 0.0 % of infected neurons export NP at 6 hpi whereas 12.1 % of infected neurons exported NP at 24 hpi.

Overall, the H7N7 NP was detected earlier in the nuclei of infected cells than the H3N2 NP, which is in agreement with the faster progression of infection (**Figure 2**). In microglia (**Figure 4B**), H7N7 showed nuclear NP localization already at 4 hpi and signs of export at 6 hpi. In H3N2-infected cells, NP localized within the cytoplasm at 6 hpi. In astrocytes (**Figure 4C**), both strains showed NP in the nucleus at 4 hpi and NP in the cytoplasm at 6 hpi. To our surprise, although NP accumulated early in the nucleus of H7N7-infected neurons (**Figure 4D**), we did not observe pronounced NP export during the entire imaging period. In contrast, the NP in H3N2-infected cells was detectable in the nuclei of infected neurons starting around 8 hpi and showed pronounced export at 16 hpi with localization to all distal parts of the cell. Taken together, as NP localization within the cytoplasm indicates nuclear export, both, astrocytes and microglia show signs of productive H7N7 infection at 6 hpi.

To further quantify the nuclear accumulation of H7N7 NP in neurons, we plotted the area of each infected nucleus together with the corresponding area of NP. At 6 hpi, we were able to differentiate two populations discriminated along the two axes (**Figure 4E**, **Supplementary Figure 5**). We interpreted the cell population with NP area larger than the nucleus (**Figure 4E**, red population) as neurons that show NP export. However, since we did not observe NP signal in the cellular periphery, this would indicate that the H7N7 NP export in neurons is limited to the soma. We further observed positive NP export with 6.1 % at 6 and 24 hpi with H7N7. In comparison, the cell population showing positive NP export after H3N2 infection was lower at 6 hpi with 2.4 % and increased to 15.9 % at 24 hpi. Taken together, as NP export is required for viral replication, especially glial cells that show high levels of infection and NP export at 6 hpi could indeed be virus producers. Regarding neurons, we detect NP export but, if productive, virus assembly would have to occur in close proximity to the nucleus. Since both, H7N7 and H3N2 show NP export in glial cells at 6hpi, but the H7N7 exerts a higher replication at 6hpi, these findings do not fully explain why the H7N7 behaves neurotropically and the H3N2 does not.

Therefore, we tested the infection and replication of both strains in the absence of microglia. In co-cultures, both strains showed lower infection rates at 6 and 24 hpi (**Figure 5A-B**) as compared to triple co-cultures. In contrast, the replication was higher in the absence of microglia for H7N7 at 6 hpi, but the differences were not statistically significant as shown in **Figure 5C**. We also did not observe differences between co- and triple co-cultured cells at 24 hpi (**Figure 5D**). To further investigate the contribution of the different cell types, we infected monocultures of astrocytes or microglia at MOI = 1 and measured the viral titer in the cell supernatants at 6 and 24 hpi with H7N7 (**Figure 5E-F**). Representative images of NP immunostainings are shown in **Supplementary Figures 8 and 9.** We observed that both cell types were able to produce H7N7 virus progeny at 6 hpi and that the titers were higher in astrocyte monocultures with 2,7×10^3^ FFU/mL (**Figure 5E**) compared to microglia monocultures with 1,1×10^3^ FFU/mL (**Figure 5F**). Since primary cultures of pure neurons were not available, we measured the titer of H7N7 in the supernatants of the infected neuronal cell line H4 at 6 hpi that showed successful infection but not replication (titer below 1,0×10^2^ FFU/mL, N=1, not shown).

**Figure 5:**
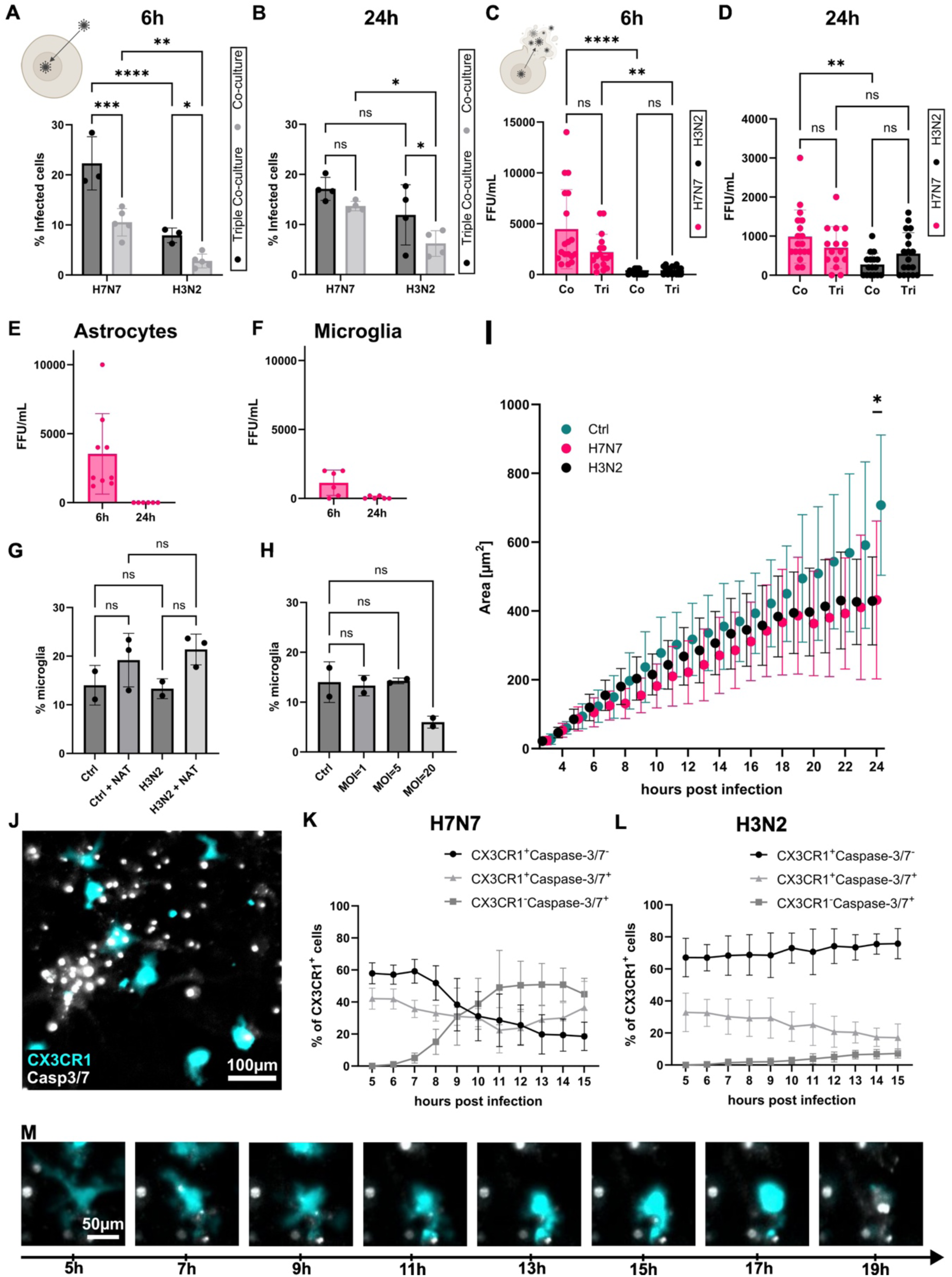
Microglia undergo apoptosis upon infection with H7N7. **A**) Infection rate of H7N7 and H3N2 at MOI=1 at 6 hpi in co- and triple co-cultures (data as in Fig. 2A). Significance calculated using two-way ANOVA followed by uncorrected Fisher’s LSD test. Co-culture with N=5; Triple co-culture with N=3. **B**) Infection rate of H7N7 and H3N2 at MOI=1 at 24 hpi in co- and triple co-cultures. Calculated using two-way ANOVA followed by uncorrected Fisher’s LSD test. N=4. **C**) Replication rate as FFU/ml of H7N7 and H3N2 at MOI=1 at 6 hpi in co- and triple co-cultures. Calculated using Kruskal-Wallis test followed by Dunn’s multiple comparisons test. N=4. **D**) Replication rate as FFU/ml of H7N7 and H3N2 at MOI=1 at 24 hpi in co- and triple co-cultures. Calculated using Kruskal-Wallis test followed by Dunn’s multiple comparisons test. N=4. **E**) Replication rate as FFU/ml of H7N7 infected astrocytic monocultures at MOI=1 at 6 hpi and 24 hpi. N=2. **F**) Replication rate as FFU/ml of H7N7 infected microglial monocultures at MOI=1 at 6 hpi and 24 hpi. N=2. **G**) Microglia concentration in cell cultures infected with H3N2 at MOI=1 in pre-or absence of externally added NAT trypsin at 24 hpi. Significance calculated using ordinary one-way ANOVA followed by Tuke’s post hoc test. N=3. **H**) Microglia concentration in cell cultures 24 h after infection with different H3N2 concentrations. Calculated using ordinary one-way ANOVA followed by Tuke’s post hoc test. N=2. **I**) Motility assay of microglia after infection with H7N7, H3N2 or control samples. Individual microglia were tracked between 5 and 15 hpi and the average Area (in pixel) was measured. Statistical analysis using two-way ANOVA with repeated measures and Tukey’s post hoc test. N=3. **J**) Representative image of triple co-cultures using CX3CR1-GFP expressing microglia (cyan) and Caspase 3/7 (Casp3/7) dye (grey). **K-L**) Caspase 3/7 activity in CX3CR1^+^ cells after H7N7 or H3N2 infection. Signal shown as % of microglia compared to 5 hpi. N=3. **M**) Panel of a single microglia that becomes Caspase-3/7 activated in the course of H7N7 infection. Statistical significances indicated as: ns: not significant, *p<0.05, **p<0.01, ***p<0.001, ****p<0.0001.

Taken together, in the presence of microglia the H7N7 infection rates increased, whereas the replication rather decreased. Experiments in monocultures showed that microglia and astrocytes were able to produce H7N7. Again, we observed a massive loss of microglia at 6 hpi with H7N7 but not H3N2.

Since IAV replication depends on the cleavage of HA0 into its fusion competent version (HA1-HA2), the presence of relevant enzymes and the cleavage success are important tropism hallmarks. While HA of H3N2 carries a monobasic cleavage site, HA of H7N7 contains a multibasic cleavage site (MCS), which might be a replication advantage in our system. We thus added external trypsin (N-Acetyl-Thiazolium-Bromid) to the cells during the infection with H3N2 to support HA activation. Indeed, externally-added trypsin had an impact on H3N2 infection and led to an increased H3N2 infection rate from 11,1 % to 17,25 % (**Supplementary Figure 7D**). However, the NAT treatment did not induce microglia disappearance after infection (**Figure 5G**). We next infected the triple co-culture with higher H3N2 concentrations and observed microglia disappearance starting only at MOI = 20 at 24 hpi, but the difference was not statistically significant (**Figure 5H**). The amount of infected cells did not show significant differences between different H3N2 concentrations at 24 hpi (F(3,4)=4.053 with P=0.1049) (**Supplementary Figure 7E**).

### H7N7 induces apoptosis in microglia

We next wanted to further investigate the response of microglia to the infection. As mentioned above, IAV replication did not decrease in the absence of microglia, although they were highly infected and showed signs of productive infection. We thus looked at microglia mobility and apoptosis during IAV infection using live-cell imaging. To better follow microglia with live-cell imaging, we used cells from CX3CR1-GFP mice that were added to Bl6 mice-derived neuronal co-cultures. When we infected these cultures, we observed a trend of a decrease in microglia mobility upon both H7N7 and H3N2 infection that was measured as the area that each living cell moves within (**Figure 5I**). H3N2-infected cells showed a statistically significant decrease in mobility compared to control samples after 24 hpi while the decrease was not statistically significant in H7N7-infected cells (P_24hpi_Ctrl-H3N2_=0.0495; P_24hpi_Ctrl-H7N7_=0.12). To study the loss of microglia during the first day of infection, we performed a live-cell Caspase-3/7 activation assay using CX3CR1-GFP expressing microglia. A representative image is shown in **Figure 5J**. Triple co-cultures were infected with H7N7 or H3N2 and microglia were tracked and subdivided based on the cellular fluorescence signal of CX3CR1 and the Caspase-3/7 marker (**Figure 5K - L**). At 15 hpi with H7N7, the composition of both signals was different compared to H3N2-infected or control samples (**Supplementary Figure 7F**). In contrast to H3N2 and controls, the amount of CX3CR1^+^/Caspase^-^ cells decreased, the amount of CX3CR1^+^/Caspase^+^ cells increased, while the amount of CX3CR1^-^ /Caspase^+^ cells also increased. In **Figure 5M**, a microglia that loses its CX3CR1 signal and becomes Caspase-3/7 positive is shown over the time course of infection. Furthermore, to understand if the Caspase-3/7 activation was ultimately followed by cell death, we visualized cell death during the course of infection. Indeed, the H7N7 infection led to an increased cell death in microglia, indicating the induction of apoptosis after Caspase-3/7 activation (**Supplementary Figure 7G and H**). Taken together, our live-cell imaging experiments showed that microglia undergo cell death by apoptosis within the first 24 hpi after infection with H7N7.

### Inflammatory profile in the CNS in vitro in the early stages of infection

IAV infections can lead to an exaggerated immune response that contributes to pathogenesis and might lead to the development of NDDs. Interestingly, non-neurotropic strains were also able to induce neurological long-term impairments [11, 24]. This clearly highlights the important role of cytokines in CNS pathologies. Therefore, we were interested in understanding early immune responses of neuronal cells once IAV enters the CNS. To this end, we took cytokine profiles upon H7N7 infection using a bead-based multiplex immune assay. In the presence of microglia, we detected several different cytokines between 6 and 24 hpi, whereas the levels of most cytokines were under the detection limit in the absence of microglia (**Figure 6A and F, Supplementary Figure 10**).

**Figure 6:**
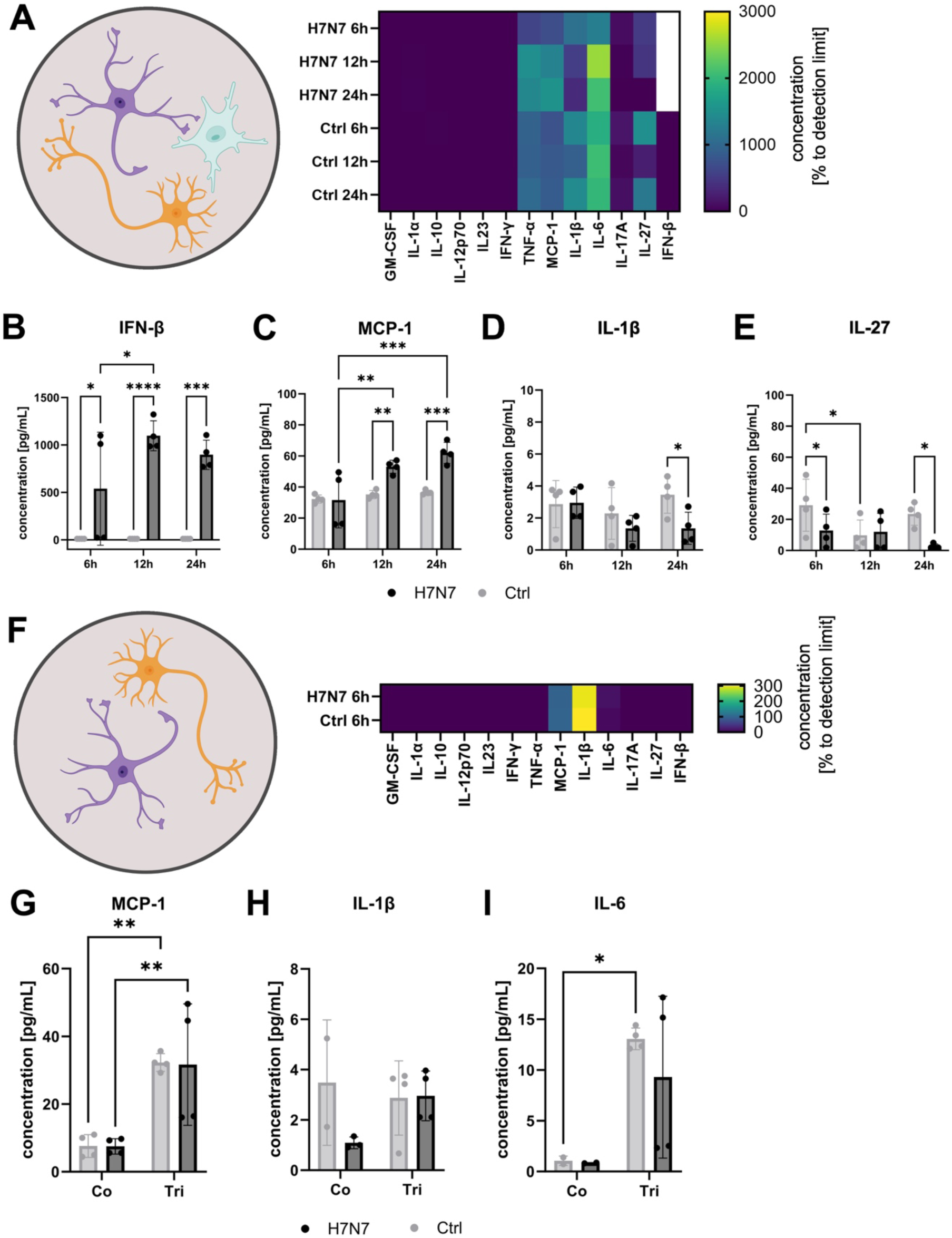
Neuroinflammatory profile after H7N7 infection. **A**) Heatmap showing relative protein levels as protein concentration compared to detection limit (in %) in the triple co-culture after H7N7 infection at MOI=1. **B-E**) Concentration of indicated cytokines compared to mock-infected control. Significance calculated using Tukey’s multiple comparison test. **F**) Heatmap showing relative protein levels as protein concentration compared to detection limit (in %) in the co-culture after H7N7 infection at MOI=1 at 6 hpi. **G-I**) Concentration of indicated cytokines in the co-culture after H7N7 infection compared to mock-infected control. Significance calculated using Fisher’s LSD test. **H**) Concentration of IL-1β compared to control calculated using Fisher’s LSD test. **I**) Concentration of IL6 compared to control. Significance calculated using Fisher’s LSD test. B-E and G-I calculated using two-way ANOVA followed by Tuke’s post hoc test. N=1. Statistical significances indicated by: *p<0.05, **p<0.01, ***p<0.001, ****p<0.0001. N=1.

The strongest effect was observed for the expression of interferon-β (IFN-β) (**Figure 6B**) in the presence of microglia, showing an increase already at 6 hpi and peaked expression at 12 hpi. IFN-β was not detectable in the absence of microglia at 6 hpi, indicating a fundamental role for microglia in the expression of this cytokine *in vitro*. Also of note, the expression of Monocyte Chemoattractant Protein-1 (MCP-1) showed an increase starting at 12 hpi compared to control samples and further increased during the course of infection (**Figure 6C**). We detected a downregulation of IL-1β at 24 hpi and Interleukin-27 (IL-27) at 6 and 24 hpi after H7N7 infection compared to control samples in the triple co-culture (**Figure 6D,E**).

In the absence of microglia, the MCP-1 level was lower for both infected and control samples at 6 hpi (**Figure 6G**). IL-1β did not show statistically significant secretion differences in the pre- and absence of microglia (**Figure 6H**). Furthermore, Interleukin-6 (IL-6) was detectable in the pre- and absence of microglia but did not show expression changes after infection (**Figure 6I**, **Supplementary Figure 10B**). Taken together, the presence of microglia was associated with elevated cytokine and chemokine expression, as early hours of H7N7 infection showed a pronounced increase in the levels of IFN-β and MCP-1.

## Discussion

In this study, we examined the infection and replication kinetics of neurotropic IAV strains in murine primary neuronal cells, along with the cellular responses to infection, to gain a deeper understanding of IAV neurotropism and its potential role in the development of NDD.

### Microglia as self-sacrificers in vitro

We observed high NP levels in microglia irrespective of the virus strain or the viral concentration. Despite microglia are macrophages, we conclude that they are indeed infected rather than phagocytosing the viruses, as the NP signal accumulates in microglial nuclei. In the presence of microglia, the overall infection rate was higher, whereas the viral replication decreased. We observed an increase in caspase-3/7 activation in microglia after H7N7 infection and, ultimately, a massive cell death. Experiments in mono-cultured microglia showed that H7N7 was able to replicate in microglia but that the titer in the supernatant was decreased in comparison to the supernatants of infected co-cultures. Together, this indicates that in early IAV infection, microglia in their function as immune cells may sacrifice themselves to protect other neuronal cell types. Our finding are in line with a previous study that observed microglia decrease following H7N7 infection *in vitro* [12]. In contrast, another recent study showed microglia increase during the acute phase of H7N7 infection *in vivo* [25]. In that study, brains from infected animals were analyzed at 8 and 10 days post-infection. Since at least some neurotropic IAV strains reach the CNS already around day 4 post infection [26], this leaves a time gap in the understanding of the initial reaction of microglia towards H7N7 reaction *in vivo*. Microglia might proliferate during the acute phase *in vivo* to present antigens to T cells, thereby enhancing the immune response and help to T cells to limit the virus infection. In our *in vitro* model microglia have to operate independently throughout the infection which *in vivo* might actually take place during the initial phase of CNS infection when T cells have not entered the CNS yet [27]. It is possible that upon H7N7 infection, microglia that attract viral particles initially show a decline in total cell numbers that may limit virus spreading and is ultimately followed by an exaggerated re-population.

Externally-added trypsin to the infection medium led to increased infection rates of H3N2, indicating an effect of the H7N7 MCS on its replication success. However, this did not induce the microglia loss and also HPAIV H5N1 did not induce the same amount of microglia disappearance as observed for the HPAIV H7N7 at lower viral titers. Therefore, the severe microglia response towards H7N7 was likely not solely attributable to its MCS. Taken together, microglia become highly infected with IAV and are able to replicate H7N7. However, since replication was higher without microglia, which die during the course of infection, neurons or astrocytes may act as main virus producers.

### Neurons exert a distinct infection profile for H7N7

IAV replication in neurons *in vitro* was shown before for several HPAIV IAV strains [28, 29]. Here, in contrast to H3N2, neurotropic H7N7 did not show NP export into the distal parts of neurons. This raises the possibility of two infection outcome scenarios in neurons: First, neurons are virus producers and viral assembly takes place at the plasma membrane around the soma. A recent study compared the sorting of the surface proteins of different viruses, including an IAV strain and observed that the glycoprotein sorting happens in a polarized manner [30]. Future studies using high-resolution microscopy techniques could shed light on any potential polarization mechanisms in the IAV replication cycle. Second, neurons are not virus producers. In this case, the NP export seems to be impaired and this disruption can happen via intra- and intercellular signals that further exert direct and indirect effects on the infected neuron. Considering that the nuclear export of vRNPs is required for viral replication and that the H7N7 NP was mostly retained in the nuclear area, we conclude that neurons likely do not contribute to virus replication.

### Astrocytes as H7N7 producers

Concluding from our results, we suggest that astrocytes are likely the main virus-producing cell type of the neurotropic H7N7 stain. This suggestion is based on three major findings: First, the H7N7 replication peaked at 6 hpi and astrocytes show pronounced nuclear NP export. Second, astrocytes had a higher contribution to the infected cell population at earlier time points compared to later stages. Third, we found high titers in the supernatant of infected astrocytic mono-cultures.

Interestingly, the cell tropism of H7N7 at 6 hpi was concentration-dependent between 4*10^6^ FFU/ml (MOI=1) and 2*10^5^ FFU/ml (MOI=0.05). The viral loads were chosen carefully based on an *in vivo* study that found titers of up to 10^5^ to 10^7^ FFU/ml of IAV in the brains of acutely infected ferrets [31]. The total infection rate increased from 10.9 % infected cells at MOI=0.05 to 22.0 % at MOI=1 corresponding to +101.8 %. Increasing the viral load in the cell culture switched the neuron-astrocyte-microglia ratio within the infected population from ∼4:3:3 at MOI=1 to ∼3:4:3 at MOI=0.05 (Neurons: MOI_0.05_=39.7 %; MOI_1_=27.0 %; Astrocytes: MOI_0.05_=31.9 %; MOI_1_=42.7 %; Microglia: MOI_0.05_=28.4 %; MOI_1_=30.4 %). By comparing the amount of cells of a particular cell type that were infected we found that this switch was mainly promoted by an immense increase of infected astrocytes at higher viral loads. While the virus load increase enhanced the amount of infected neurons and microglia by +17.6 % and +42.3 %, respectively (Neurons: MOI_0.05_=11.9 %; MOI_1_=14.0 %; Microglia: MOI_0.05_=40.0 %; MOI_1_=56.9 %) it increased the amount of infected astrocytes by even +233.3% (MOI_0.05_=6.0 %; MOI_1_=20.0 %). It is reasonable to assume a mechanism in which glia cells protect the precious neurons. The high amount of infected microglia after infection with the neurotropic and the non-neurotropic IAV strain, a potential restriction of H3N2 especially at lower viral titers as well as the induced cell death after H7N7 infection indicate microglia act as an initial IAV barrier. It was shown that a microglial barrier restricted the spreading of vesicular stomatitis virus into the CNS via the olfactory bulb and that this mechanism was dependent on IFN-β signaling [32]. Our findings further support a mechanism, in which astrocytes step in as a second barrier if microglia protection is not sufficient, for example as observed at high IAV titers here.

Our findings align with an *in vitro* study that found the replication in astrocytes to be restricted to neurotropic IAV variants [29]. Furthermore, this study observed the MCS and to a lesser extent the viral attachment to be an important virulence factor. Here, astrocytes did not show GlcNAc or Neu5Ac sialic acids on their surface shown by WGA, SNA and MAA1 staining [33]. As H7N7 was able to enter astrocytes this suggests both IAV strains utilize another primary attachment factor in astrocytes.

### Inflammatory potential of different IAV strains

Both, neurotropic and non-neurotropic H7N7 and H3N2 induced cognitive long-term impairments in infected mice well beyond the acute phase of infection *in vivo* [11]. H3N2 was tested in a transgenic mouse model (APP/PS1) and was able to induce AD symptoms *in vivo* [24]. Furthermore, H7N7 infection was shown to exert sex-specific differences in neurons, astrocytes and microglia *in vitro* [12]. As NDDs have a strong socio-economic impact [34, 35], prevention is of great interest. A study by Levine et al. identified several viral exposures that were associated with NDD and among them especially viral encephalitis and Alzheimer’s disease had the largest effect association [3]. Furthermore, neurotropic IAV was able to induce prion protein misfolding *in vitro* [36] [37]. This indicates that neurotropic IAV strains such as H7N7 have an even higher impact on the onset and progression of NDD.

We performed a cytokine assay to identify the neuroinflammatory profile upon H7N7 infection.

Our data indicates that microglia are the primary producers of inflammatory mediators post-infection with H7N7, but are dying beyond 6 hpi. Most cytokines and chemokines show half-times ranging between minutes (TNF-α, MCP-1, IL-27 and IL-1β) to hours (IFN-β, IL-6) [38–41]. Focussing on the timing of cytokine and chemokine production following infection we found molecules such as IFN-β which is critical for immediate viral suppression, and MCP-1 which is involved in the recruitment of peripheral immune cells [42, 43] represent the early-stage immune response. Neurons, microglia and astrocytes can produce MCP-1 [44, 45]. Microglia mobility was impacted after infection with IAV. The largest effect was observed for the expression of IFN-β. This cytokine is of “Dual Nature”: While the acute production of IFN-β exerts a beneficial antiviral activity, the chronic expression induces a disadvantageous inflammatory state [46]. We found, after H7N7 infection, there was a severe upregulation of IFN-β. Moreover, we found IL-27 that exerts an antiviral function via STAT1/2/3 and PKR phosphorylation [47] was downregulated during the early phase of infection. As *in vivo* studies observed an upregulation [47, 48], this indicates cell reactions may reverse during the course of infection and that there is a lack of peripheral immune cues in our *in vitro* model.

Cytokines like TNF-α, IL-1β and IL-6 did not show changes in the protein levels within the first 12 hpi and might therefore represent the later stage in the immune response. These cytokines were reported to be involved in processes associated with PD and AD pathogenesis [49–51]. We found a downregulation of IL-1β at 24hpi with H7N7. Since microglia had already disappeared at 24 hpi the detection of cytokines involved in the later stage of infection was more unlikely compared to cytokines and chemokines involved in the very early stages of infection. Taken together, we observed a severe inflammatory response towards H7N7 infection in neuronal cells *in vitro*. Future studies should include further neurotropic as well as non-neurotropic variants.

We tested the infection rate of the avian H5N1 clade 2.3.4.4b (A/black-headed gull/Ger-BW/AI01419/2023). Despite not being mouse-adapted, this IAV strain induced a robust infection rate in neuronal cells that was even higher compared to mouse-adapted H3N2 that behaves non-neurotropic *in vivo* [11]. H5N1 IAVs exert a broad neurotropic potential [6]. Therefore, both HPAIV were able to induce a robust infection whereas the non-neurotropic H3N2 was not ([11], reviewed in [6]). *In vitro* studies by Siegers et al. found that H5N1 replication depends on its MCS and to a lesser extent on cell attachment. Moreover, they showed that in contrast to H3N2 that did show replication only in a trypsin-dependent manner in neurons, the replication of H5N1 was possible in neurons and astrocytes [29]. However, it is not clear whether H7N7 and H5N1 use the same mechanisms.

Taken together, we observed that neurotropic H7N7 induced a robust infection in triple co-cultured neurons, astrocytes and microglia and triggered a massive microglia cell death accompanied by an exaggerated level of IFN-β and other cytokines. We suggest that neurotropic H7N7 can replicate in astrocytes and microglia and to a lesser extend in neurons as well. Our data indicate that of all cell types, particularly the replication in astrocytes was most beneficial for H7N7 replication.

## Acknowledgements

We would like to thank Jennifer Fricke, Diane Mundil and Tina Geisler for excellent technical assistance. This work was supported by the Helmholtz Association (VH-NG-1526 to C.S.). C.S. and M.K. acknowledge support through the cooperativity and creativity project call (CCC, project 3) of HZI.

