## Supplementary Material for "Glial cells promote infection by neurotropic Influenza A viruses *in vitro*"

Supplementary Table 1: List of antibodies

| Name | Company | Product Number | Dilution |
|---|---|---|---|
| Mouse-α-Nucleoprotein | Merck | MAB8257 | 1:500 |
| Rabbit-α-H7N7 HA | Invitrogen | | 1:500 |
| Rabbit-α-H3N2 HA | Invitrogen | PA5-34930 | 1:500 |
| Chicken-α-NeuN | Synaptic Systems | 266006 | 1:500 |
| Guinea pig-α-MAPII | Synaptic Systems | 188004 | 1:500 |
| Guinea pig-α-GFAP | Synaptic Systems | 173004 | 1:500 |
| Rabbit-α-GFAP | Invitrogen | PA1-10019 | 1:500 |
| Guinea pig-α-Iba1 | Synaptic Systems | 234308 | 1:500 |
| Goat-α-Guinea pig conjugated with Alexa Fluor 488 | Invitrogen | A-11073 | 1:500 |
| Goat-α-Guinea pig conjugated with Alexa Fluor 647 | Invitrogen | A-21450 | 1:500 |
| Goat-α-Chicken conjugated with Alexa Fluor 488 | Invitrogen | A-11039 | 1:500 |
| Goat-α-Chicken conjugated with Alexa Fluor 647 | Invitrogen | A-21449 | 1:500 |
| Goat-α-Mouse conjugated with Alexa Fluor 568 | Invitrogen | A-11031 | 1:500 |
| Goat-α-Rabbit conjugated with Alexa Fluor 568 | Invitrogen | | 1:500 |
| Wheat germ agglutinin conjugated with Alexa Fluor 647 | Invitrogen | W32466 | 1:500 |
| Sambucus Nigra Lectin conjugated with FITC | Vector Laboratories | FL-1301-2 | 1:200 |
| Maackia Amurensis conjugated with Fluorescin | Vector Laboratories | VEC-FL-1311 | 1:200 |

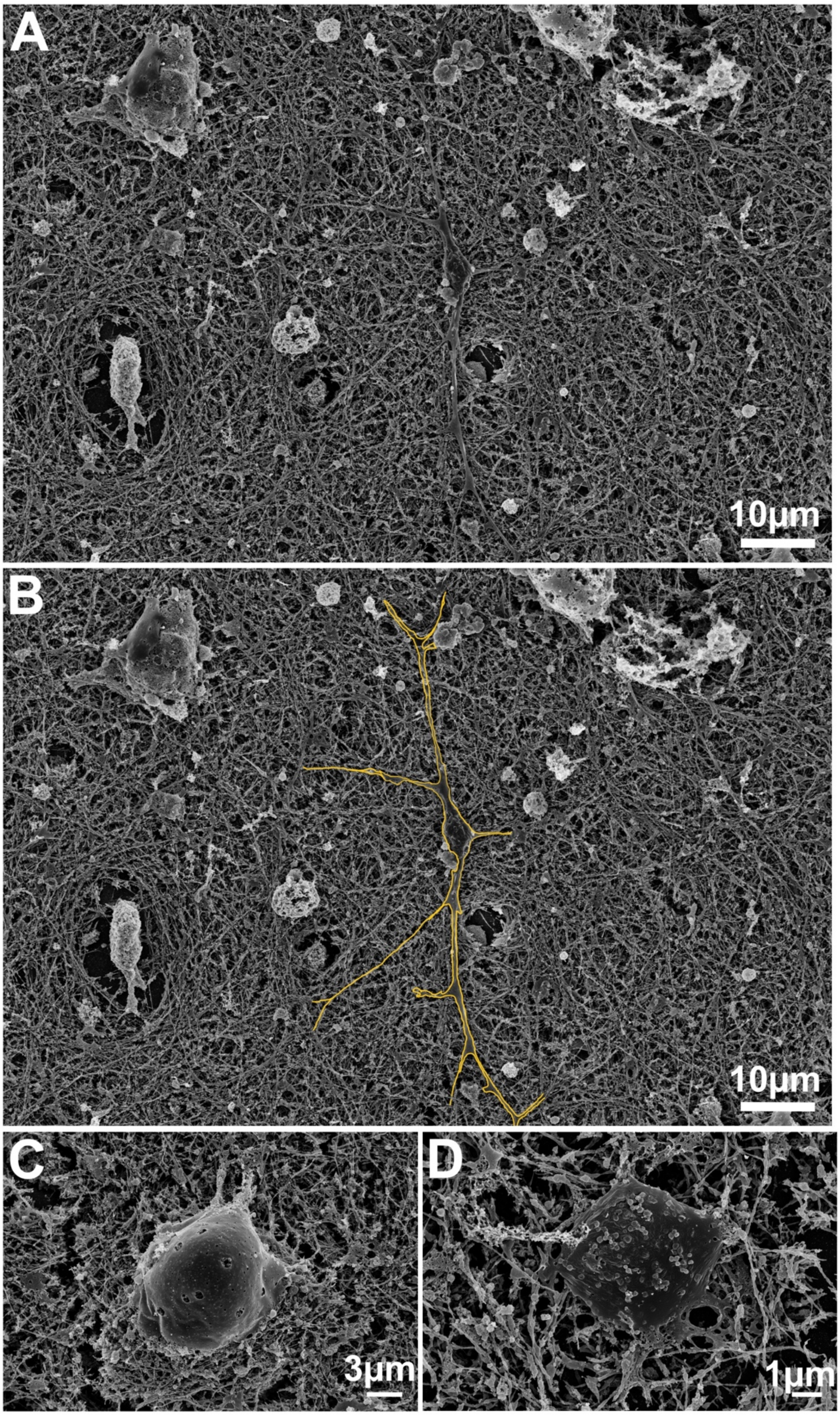


**Supplementary Figure 1:** Scanning Electron Microscopy (SEM) of murine primary CNS triple co-culture. **A-B**) Overview images demonstrating the dense cellular network. **C**) Cell surface of a cell in the control sample. **D**) Cell surface of an infected cell.

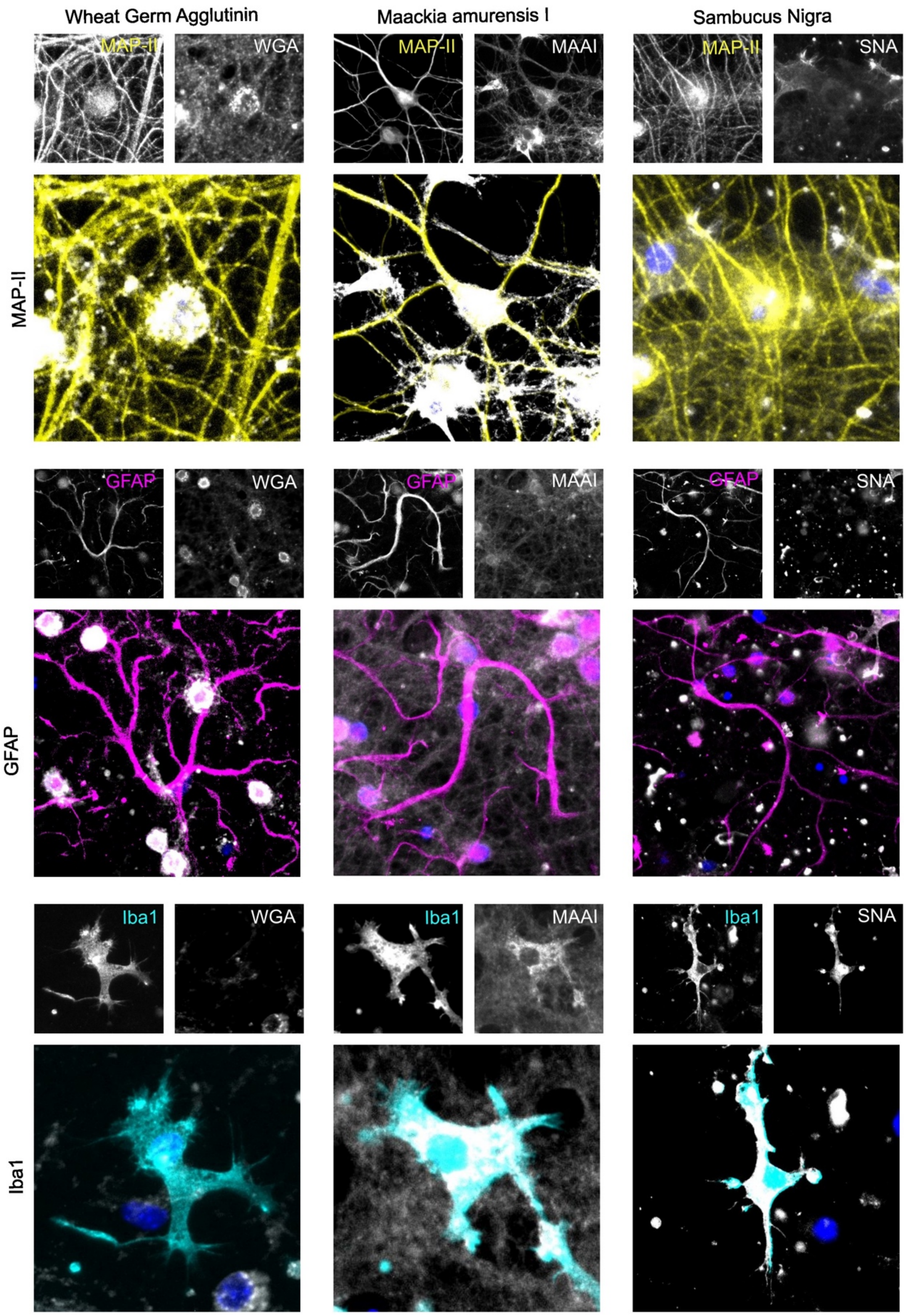


**Supplementary Figure 2:** Lectin staining of murine primary CNS triple co-culture. Yellow indicates neurons (MAP-II), purple astrocytes (GFAP) and cyan microglia (Iba1). Wheat germ agglutinin, Maackia amurensis I and Sambuccus nigra are shown in grey scale.

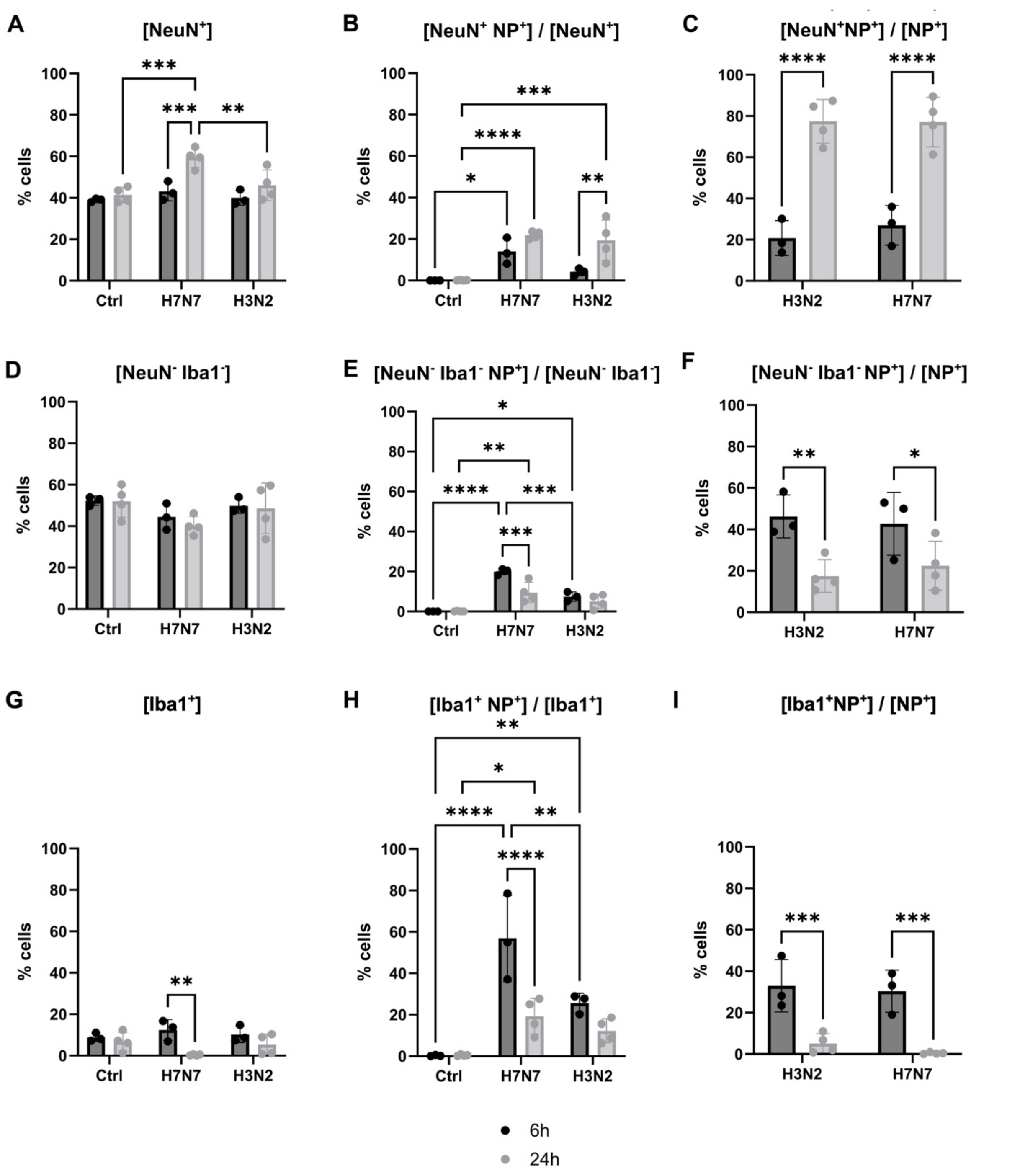


**Supplementary Figure 3:** Cell type-specific infection rates at MOI=1. **A-C**) neurons, **D-F**) astrocytes and **G-I**) microglia. **A, D, G**) Total cell population. **B, E, H**) Percent of cells infected for each cell type. **C, F, I**) Percent cells of a particular cell type within the infected cell population. A, B, D, E, G, H) Significance calculated using two-way ANOVA followed by Tukey's post hoc test. C, F, I) Significance calculated using two-way ANOVA followed by uncorrected Fisher's LSD test. N=5, n<1. Statistical significances indicated by: *p<0.05, **p<0.01, ***p<0,001, ****p<0,0001. 6 hpi: N=3; 24 hpi: N=4.

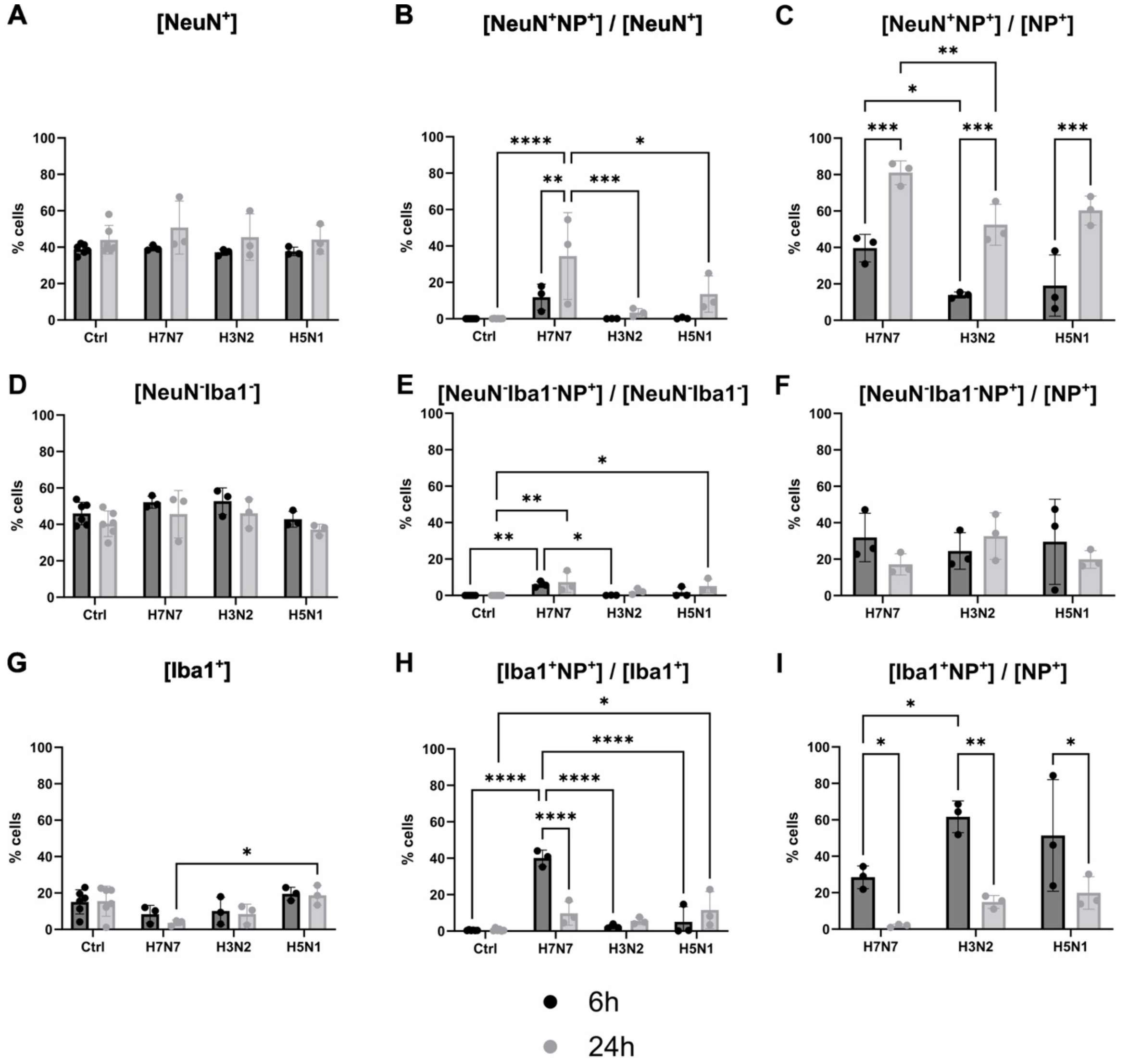


**Supplementary Figure 4:** Cell-type specific infection rates at MOI=0.05. **A-C**) neurons, **D-F**) astrocytes and **G-I**) microglia. **A, D, G**) Total cell populations. **B, E, H**) Percentage of cells infected for each cell type. **C, F, I**) Percent cells of a particular cell type within the infected cell population. Significance calculated using two-way ANOVA followed by Tukey's post hoc test. N=5. Statistical significances indicated by: *p<0.05, **p<0.01, ***p<0,001, ****p<0,0001. N=5; n=1.

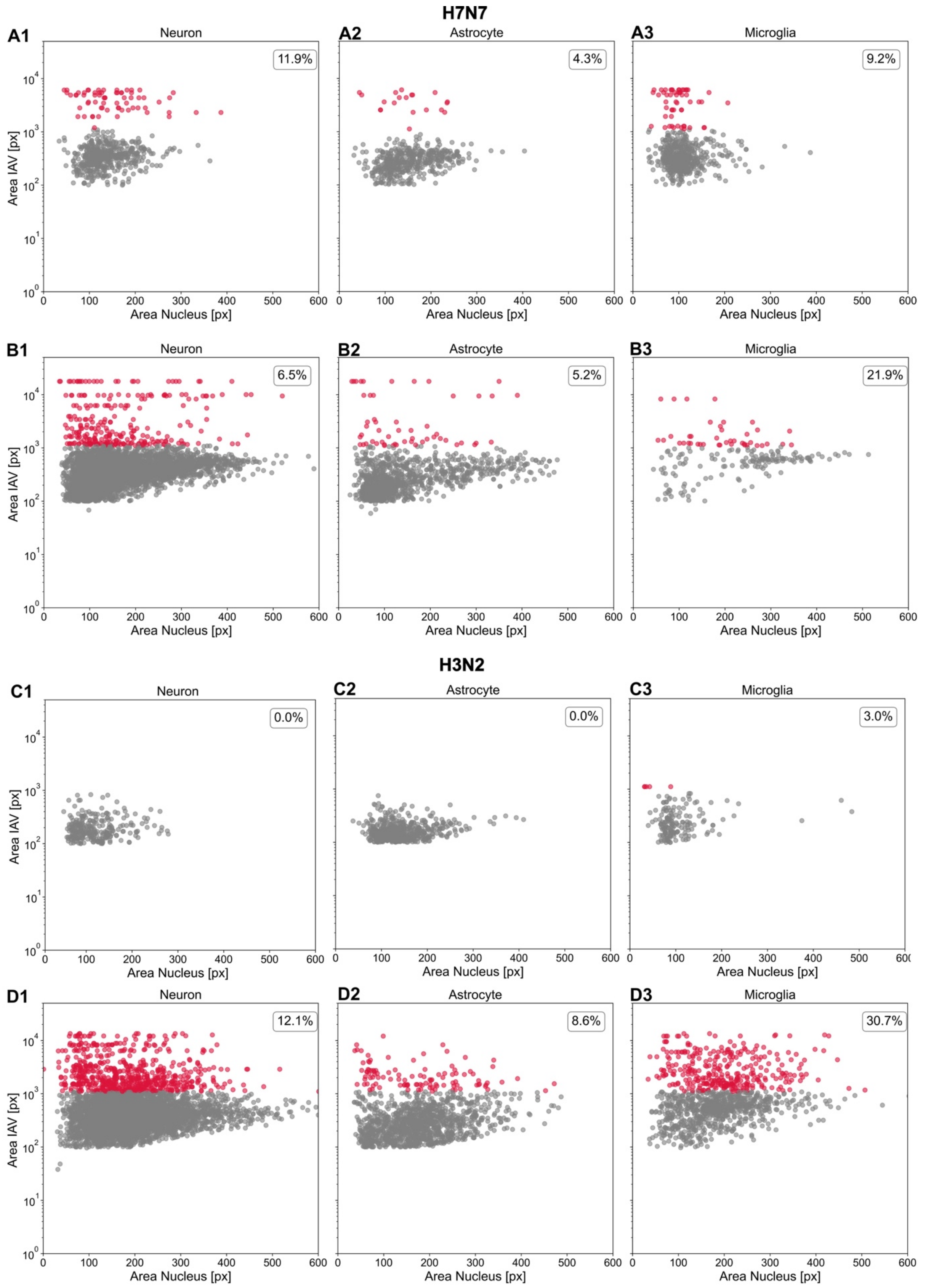


**Supplementary Figure 5:** Percentage of NP export per cell type after infection with H7N7 (panel A-B) or H3N2 (panel C-D) at MOI=1. Upper panels (A and C) show 6 hpi and lower panels show 24 hpi (B and D). N=5.

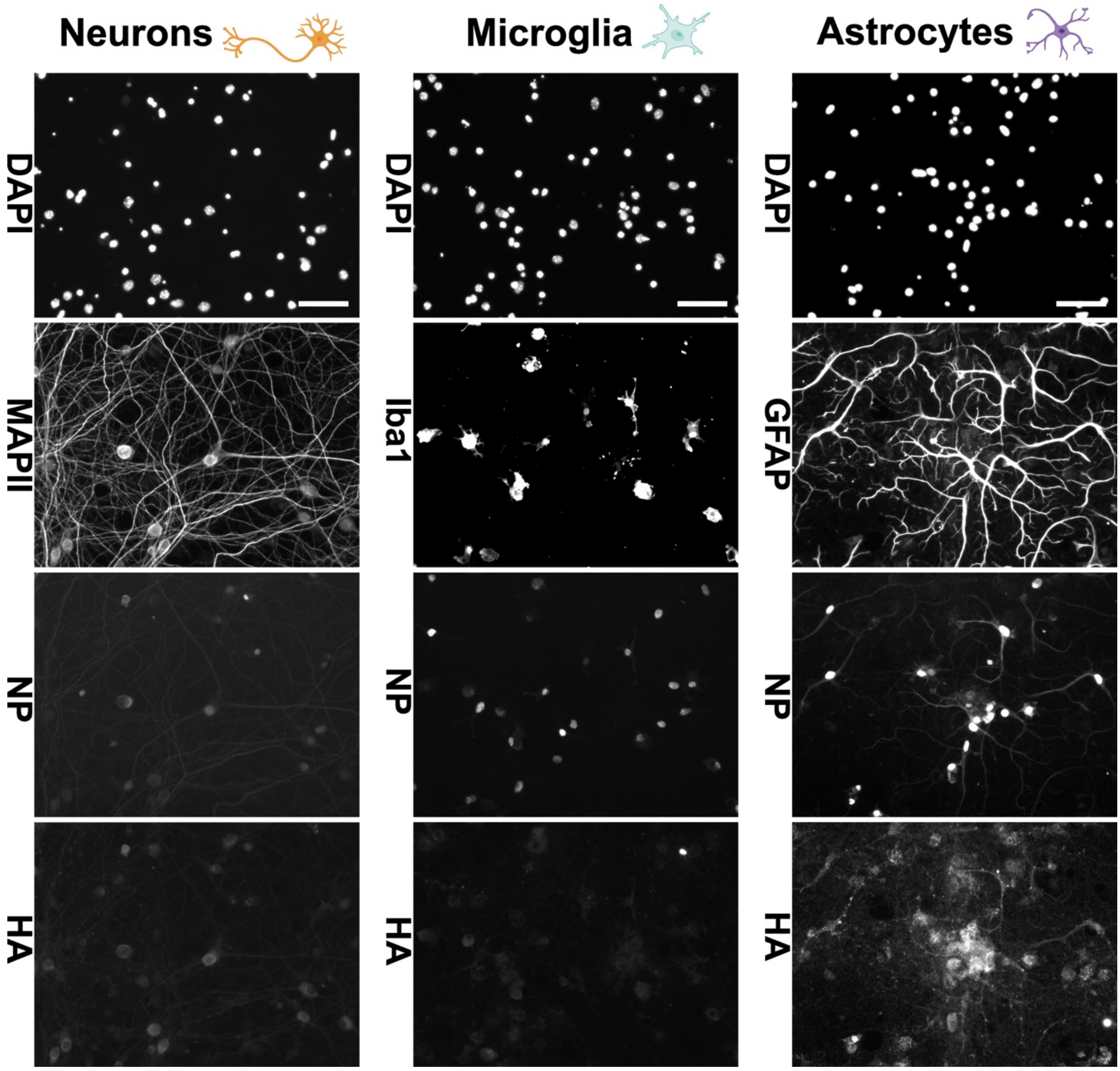


**Supplementary Figure 6**: Overview images of neurons (left), microglia (middle) and astrocytes (right) after H7N7 infection, stained for DAPI (first row), cell markers (second row), NP (third row) and HA (last row). Scale 50µm.

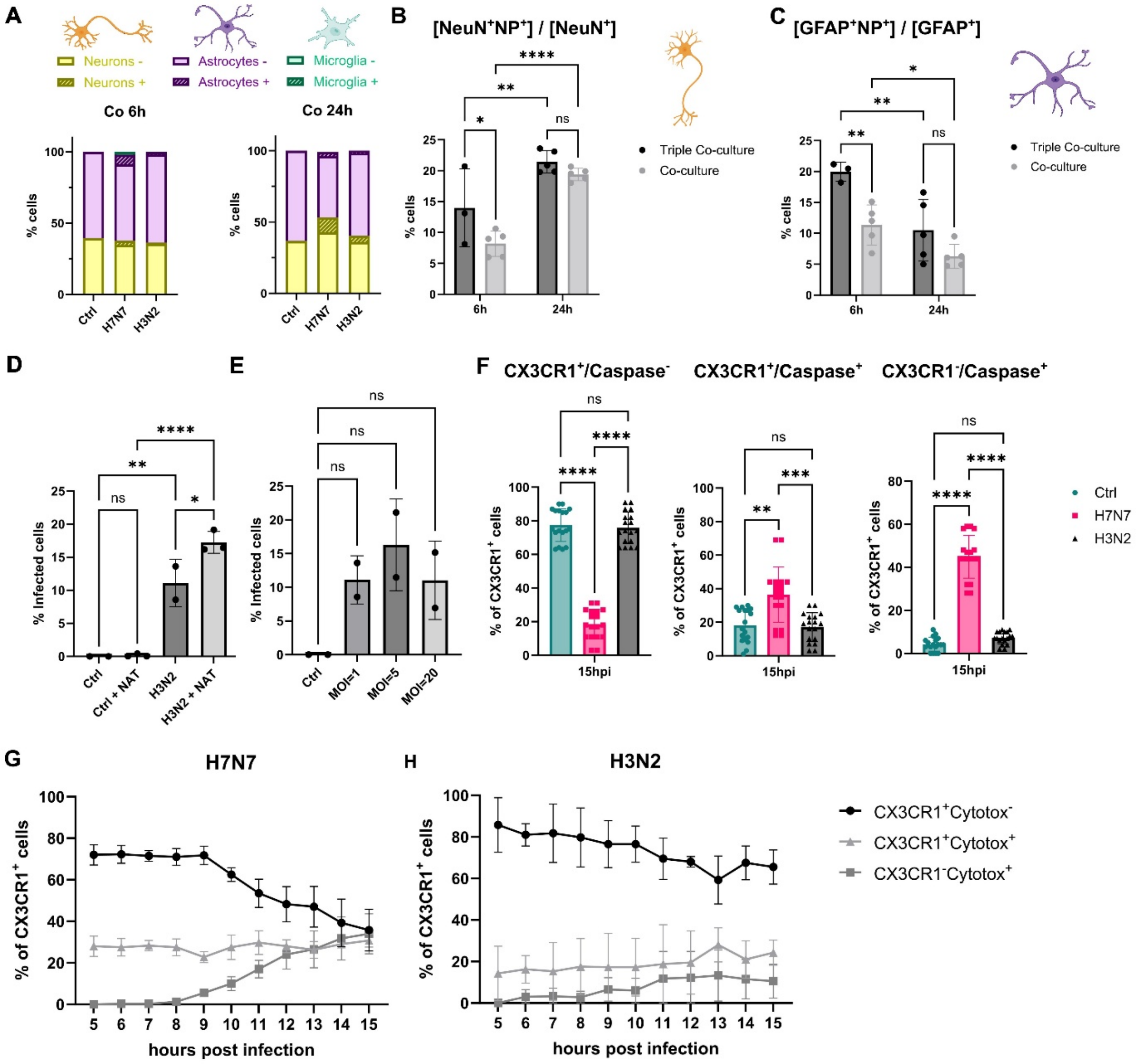


**Supplementary Figure 7: A**) Cell culture composition of co-cultures at 6 and 24 hpi. Dashed areas indicate infected cells. **B**) Percent of infected neurons in co- or triple co-cultures at MOI=1. N=5. **C**) Percent of infected astrocytes in co- or triple co-cultures at MOI=1. N=5. **B-C** Significance calculated using two-way ANOVA followed by uncorrected Fisher's LSD test. **D**) Infection rate of H3N2 in the pre- or absence of externally added trypsin at 24 hpi. N=3. **E**) Infection rate of H3N2 at different MOIs at 24 hpi. D-E Significance calculated using two-way ANOVA followed by Tukey's post hoc test. N=2. **F**) Microglia subpopulations at 15 hpi based on the presence of CX3CR1 and caspase 3/7 marker. Calculated using Kruskal-Wallis test followed by Dunn's multiple comparisons test. N=3. **G-H**) Cell death analysis. Different microglia (CX3CR1) subpopulations are analyzed in the course of H7N7 (G) or H3N2 (H)

infection based on the presence of CX3CR1 and cell death marker. N=1. Statistical significances indicated by: ns: not significant, *p<0.05, **p<0.01, ***p<0,001, ****p<0,0001.

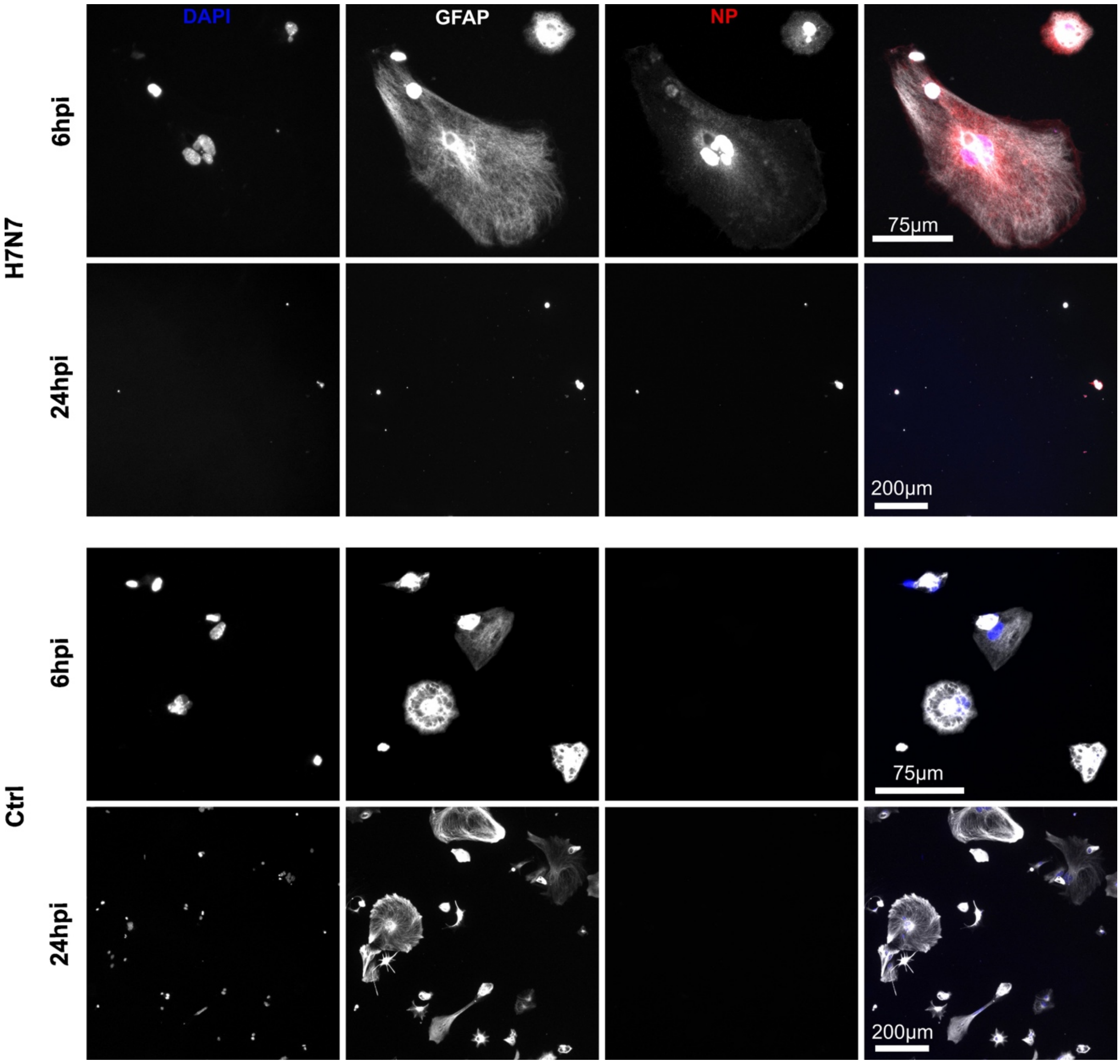


**Supplementary Figure 8:** Infection of astrocytic monoculture with H7N7 or PBS as control for 6 and 24 hpi.

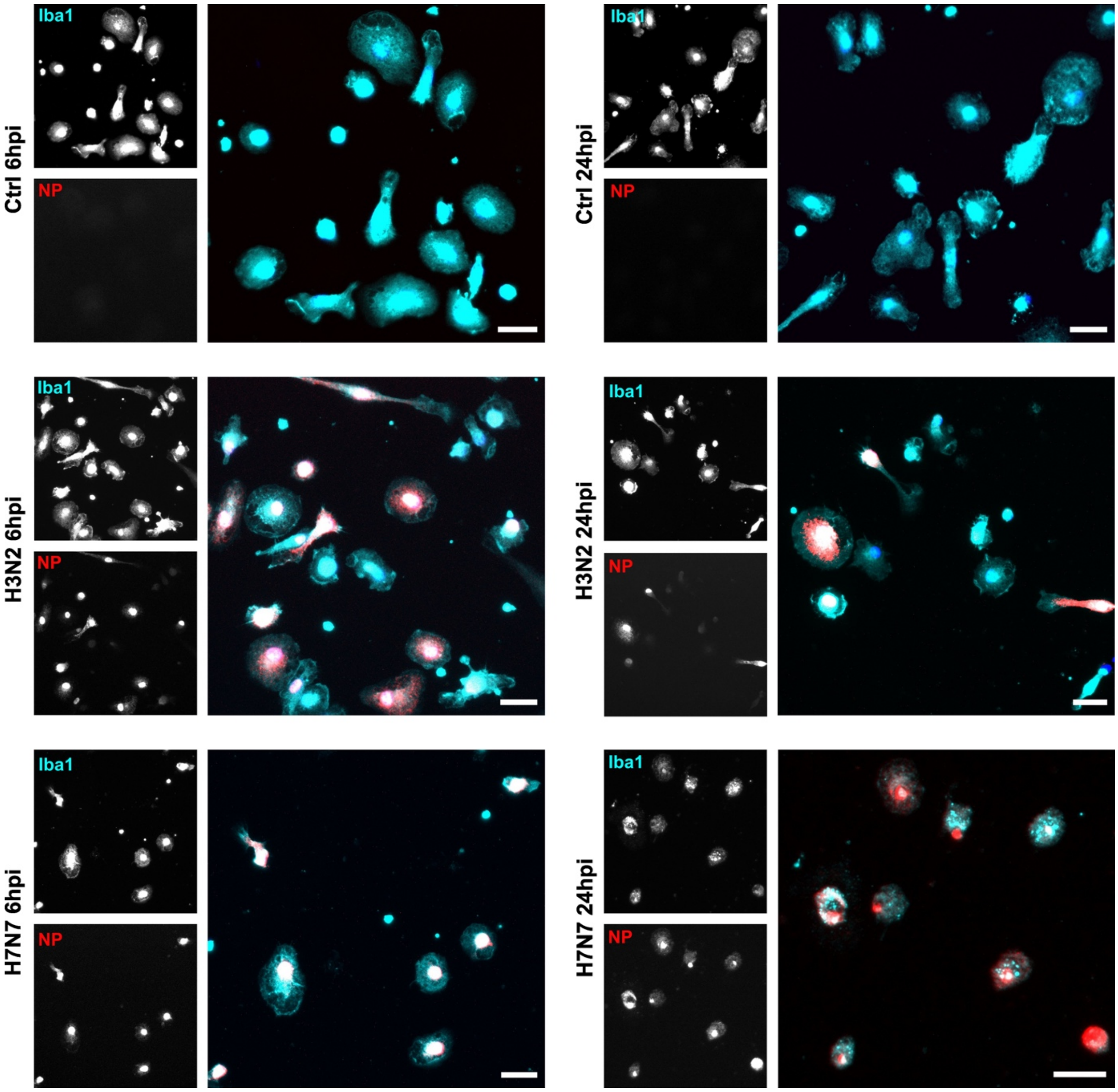


**Supplementary Figure 9:** Infection of microglia monocultures with H7N7 or PBS as control for 6 and 24 hpi stained for Iba1 and NP. Scale: 20µm.

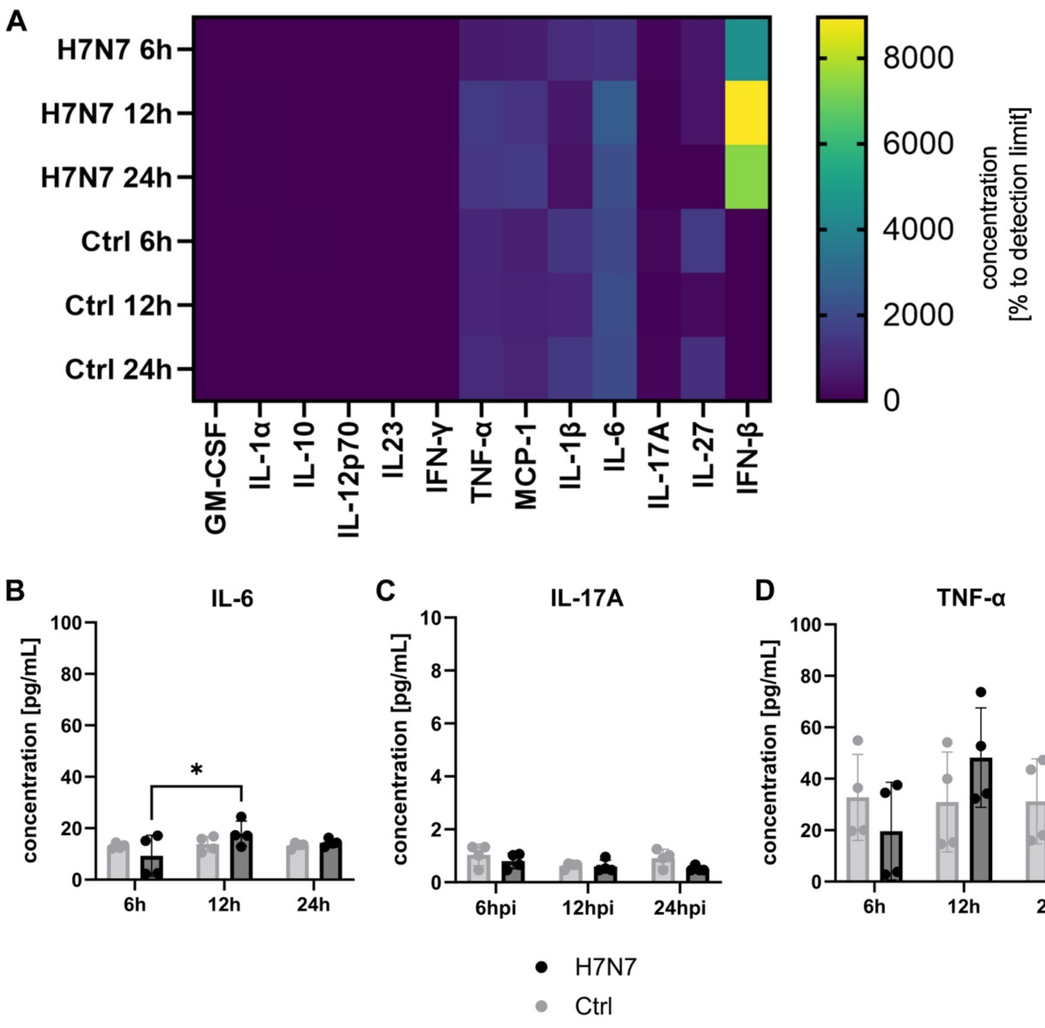


**Supplementary Figure 10:** Cytokine screening after H7N7 infection. **A**) Heatmap showing cytokine concentrations as % to detection limit. **B**) IL-6 concentration at different time points between H7N7 infected cells and control samples. **C**) IL-17A concentration at different time points between H7N7 infected cells and control samples. **D**) TNF-α concentration at different time points between H7N7 infected cells and control samples. Calculated using two-way ANOVA followed by Tukey's multiple comparisons test. Statistical significances indicated by: *p<0.05, **p<0.01, ***p<0,001, ****p<0,0001. N=1, n=4.
